# Validation of a multiscale Hill-type actuator against comprehensive benchmarks of motor unit and muscle force measurements

**DOI:** 10.64898/2026.06.15.732276

**Authors:** Andrea Sgarzi, Arnault H. Caillet, Matthew Millard, Sven Weidner, Nicos Haralabidis, Théo Meranger, Bart Bolsterlee, Dario Farina, Nigel H. Lovell, Luca Modenese

## Abstract

Computational Hill-type muscle models are widely used to simulate muscle force production because of their efficiency and physiological interpretability. However, their formulation relies on limiting assumptions, including debated multiscale simplifications, a simplified excitation-activation dynamics and an inability to capture slow and fast fibres. Moreover, existing Hill-type models remain insufficiently validated across physiological scales, fibre types, and contraction modes. We addressed these limitations by developing a multiscale fibre-type specific Hill-type neuromuscular actuator with mechanistic excitation-activation dynamics and systematically validated it against comprehensive experimental benchmarks. The model built upon a previously proposed motoneuron-driven actuator incorporating calcium-kinetics-based activation dynamics. The excitation-activation formulation was further refined to strengthen its physiological basis, while the contraction dynamics was extended by including an activation- and length-dependent force-velocity relationship, elastic tendon, passive elastic element, and the fibre-type-specific effects of yielding and sag. Validation was performed against four benchmark datasets spanning motor-unit and whole-muscle scales, including slow and fast fibres under both isometric and dynamic conditions. Experimental force traces were obtained from six muscles of rats and cats using a broad range of stimulation frequencies, muscle lengths, and imposed length changes, combining previous literature datasets with experiments performed *ad hoc* for this study. Overall, the model reproduced forces across all benchmark conditions, with mean absolute errors typically below 15% of the maximum isometric force, although larger errors were observed in specific submaximal and dynamic trials. The inclusion of physiologically based excitation-activation dynamics, together with yielding and sag, improved model performance under submaximal activation conditions. This study presents the first systematic validation of a single multiscale Hill-type neuromuscular actuator against comprehensive experimental motor unit and muscle force data, providing a benchmark framework for the development and assessment of future models.

**Author summary:** Skeletal muscles generate force through a complex sequence of events that links neural signals to muscle contraction. Because direct measurements are difficult to obtain, researchers often rely on computer models to investigate neuromuscular function and estimate muscle forces. However, most modelling approaches rely on simplifying assumptions about how force is generated across different biological scales, how muscles are activated, and how slow and fast muscle fibres behave. Moreover, they have not been validated against comprehensive experimental data. As a result, it remains unclear how accurately these models can reproduce muscle force across different physiological conditions.

In this study, we established the first comprehensive set of experimental benchmarks spanning both motor-unit and whole-muscle scales, including slow and fast muscles under isometric and dynamic conditions. We used these benchmarks to validate a newly developed multiscale muscle model that explicitly represents the physiological pathway from neural stimulation to force production. The model incorporates experimentally based descriptions of calcium dynamics, activation, tendon elasticity, and fibre-type-specific contractile properties. We then compared simulated and experimental force responses across a wide range of stimulation frequencies, muscle lengths, and length-change conditions.

## 1 Introduction

Skeletal muscles generate movement and force through the coordinated activity of motor units (MUs), the functional units of the neuromuscular system [1]. Each MU consists of a single spinal motoneuron (MN) and the muscle fibres it innervates, transforming an axonal action potential (AP), the electrical command sent by the central nervous system, into an electrical and mechanical output through excitation-contraction coupling [2]. The AP propagates along the MN axon and determines the fibre depolarisation. This triggers the release of calcium ions in the sarcoplasm of the muscle fibres, initiating cross-bridge cycling and producing tension that is transmitted through the tendon to generate movement. The complexity of this process is exemplified by the multiple physiological scales involved (sarcomere, single fibre, MU and whole muscle) as well as by the diversity of skeletal muscles, which differ in their contractile properties according to fibre composition. Skeletal muscles are typically classified as slow or fast, depending on the predominance of slow or fast fibres expressing distinct myosin heavy chain isoforms [3], while MUs have been functionally divided into slow, fast fatigue-resistant, or fast fatigable based on their contractile and fatigue properties [4, 5].

Computational models are widely used to estimate the muscle forces generated by these neuromechanical processes, as experimental measurements are difficult or impossible to perform non-invasively [6]. Among these, Hill-type muscle models [7, 8] remain the most widely used due to their computational efficiency and physiological interpretability. In their most common formulation, they represent the muscle-tendon unit using a Contractile Element (CE) characterised by force-length (FL) and force-velocity (FV) relationships, a Parallel-Elastic Element (PE) and a Series-Elastic Element (SE) [9]. Despite their success in large-scale simulations of human movement [10], they present several limitations. A first fundamental issue lies in their multiscale use. Although studies such as [11] have shown that macroscopic muscle behaviour can be inferred from sarcomere-level properties, this assumption remains debated and it is still unclear whether Hill-type models can accurately reproduce force-generation mechanisms across both MU and muscle levels [9]. Secondly, the mathematical description of the muscle’s neuromechanical interface is typically reduced to a single “active state” variable [9], which cannot be directly measured or validated experimentally [12]. By assuming that the active state acts independently from the contractile properties, Hill-type models may fail to represent the nonlinear interplay between activation, stimulation frequency (*f* ^*s*^) and muscle length, especially under submaximal and dynamic conditions [9, 13-15]. Third, most Hill-type actuators do not account for the distinct neuromechanical properties of slow and fast fibres [9], although accounting for these differences has been shown to improve estimates of whole-muscle force under both isometric [16] and dynamic [17] conditions.

To address these limitations, some studies scaled the Hill-type formulation down to the individual MU level, computing total muscle force as the sum of the independent mechanical contributions of individual MUs [18-21]. Moreover, some neural impulse-driven approaches explicitly modelled additional physiological steps preceding the active state, including sarcoplasmic calcium dynamics, capturing the nonlinear dependencies on *f* ^*s*^ and muscle length [18, 20, 22], with [18-20] also distinguishing between slow and fast fibres. Nevertheless, these studies lack validation against comprehensive datasets of varying experimental conditions (i.e., varying stimulation frequencies, lengths and velocities for different fibre types), highlighting the critical need to establish benchmark simulations to systematically evaluate and compare their predictive accuracy and reusability [9, 13, 15]. However, constructing such benchmarks remains challenging due to experimental constraints. The force of single muscles [16] or MUs [1] is typically measured in animal preparations under controlled physiological conditions. Even in these cases, previous validation studies have generally been limited in scope, focusing predominantly on whole-muscle behaviour [14-16, 22-25] rather than MU-level dynamics, and on slow muscles [14, 15, 22-24, 26].

In this context, the present study introduces a fibre-type-specific Hill-type model based on the MN-driven framework of [18] and aims to validate it under isometric and dynamic conditions at both the MU and muscle scales. The model explicitly represents the excitation-contraction coupling through a set of coupled ordinary differential equations (ODEs) describing the transformations from MN discharge input to calcium release, activation, and mechanical force generation. To evaluate its physiological accuracy, we established a set of benchmark datasets across scales using both data from literature and new animal experiments performed *ad hoc* for this study. These benchmarks include data from two slow (cat soleus and rat soleus) and two fast (cat caudofemoralis and rat extensor digitorum longus) muscles, covering isometric and dynamic contractions under both maximal and sub-maximal stimulation conditions, as well as a comprehensive set of MUs from the cat and rat medial and lateral gastrocnemius under isometric conditions. By comparing simulated and experimental forces across these benchmarks, this study provides the first systematic validation of a single Hill-type neuromuscular actuator across contraction modes, fibre types, and physiological scales.

## 2 Methods

### 2.1 Hill-type model

The Hill-type model considered in this study builds upon the MN-driven framework developed by [18]. In its original formulation, the model consisted of a population of n-in-parallel Hill-type force generators (FGs), with the *i*^*th*^ FG representing a MU. Each MU transformed its corresponding stimulation input *s*_*i*_(*t*) from experimental discharge times into the active state a_*i*_(*t*) and the force 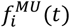, differentiating between slow and fast MU types. Each FG comprised a Neuromechanical Element (NE) modelling the excitation-activation dynamics, and a CE with constant length *l*^*CE*^ modelling the contraction dynamics. The forces 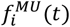 were non-linearly summed to generate the total muscle force *F*^*M*^(*t*) during isometric contractions. In this study, the single MU structure of [18], with a refined excitation-activation dynamics, has been extended to represent a fibre-type-specific neuromuscular actuator at multiple scales (MU and whole muscle) and simulate also dynamic contractions (Fig 1). Muscle properties at all scales were normalised (e.g.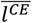) to the muscle maximum isometric force *F*_0_, optimal fibre length 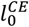 and maximum contraction velocity *v*_*max*_ as appropriate [8]. All model constitutive equations and parameters are available in the Supplementary Materials (S1-S3), with the key aspects presented in the following sections.

**Fig 1.**
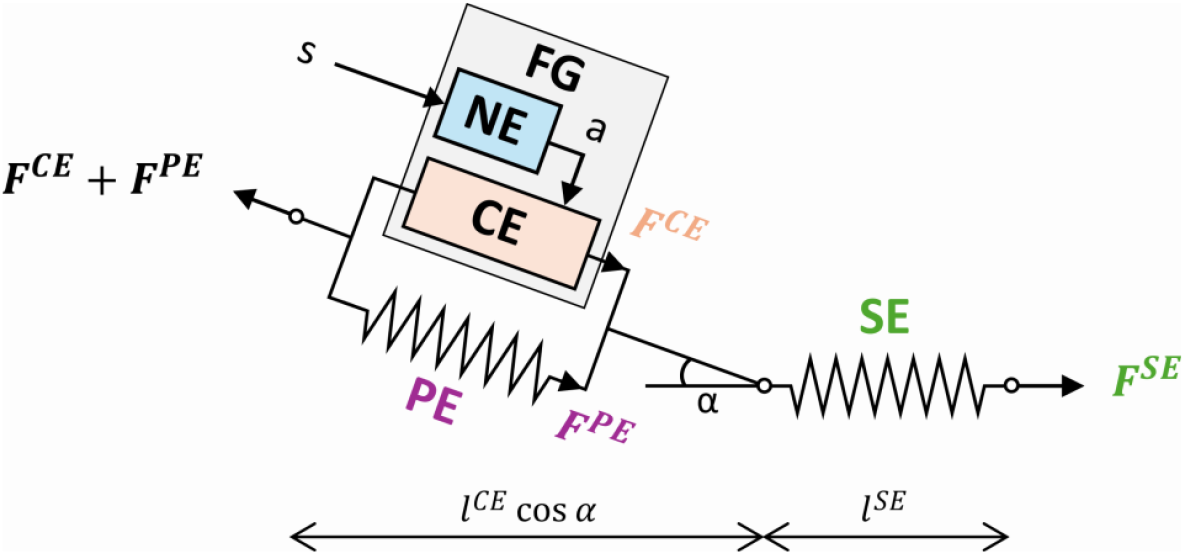
Hill-type actuator used to represent force generation at both the single motor-unit (MU) and whole-muscle scales. The model’s active force generation component is represented with a Force Generator (FG) comprising a Neuromechanical Element (NE) and a Contractile Element (CE). The NE governs the excitation-activation dynamics, transforming the neural excitation input *s* into the active state a. The active state subsequently scales the force production of the CE, which is arranged in parallel with a Parallel-Elastic Element (PE), both with length *l*^*CE*^, accounting for the contraction dynamics and passive connective tissues, respectively. A Series-Elastic Element (SE) of length *l*^*SE*^ is included to represent the elastic properties of the tendon. The architecture also incorporates the pennation angle α, defined between the parallel CE-PE components and the SE.

#### 2.1.1 Excitation and Activation Dynamics

The excitation dynamics of the proposed model transforms binary input trains, representing the MN discharge times, into APs *u*(*t*) propagating along the muscle fibres using the same equations proposed in [18], which were based on [27]. In the activation dynamics, *u*(*t*) controls the time-varying level of sarcoplasmic calcium concentration [Ca^2+^] *c*(*t*), which is described by a 2^nd^ order ODE (Eq. 1), modified from [18] as follows:

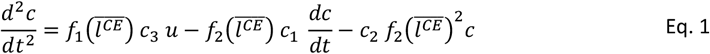

In Eq. 1, 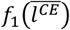 and 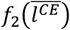 are functions that have been fit to the experimental data from [28, 29], capturing the non-linear dependence of [Ca^2+^] on 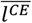 observed *in vitro* in frog muscle fibres. The fibre-type-specific coefficients *c*_1_, *c*_2_ and *c*_3_ were obtained by separately replicating the experimental conditions (sarcomere length and *f* ^*s*^) of mouse slow [30, 31] (Fig 2A) and fast fibres [32] (Fig 2B) whilst minimising the squared error between simulated and measured [Ca^2+^] levels. Additional methodological details, including considerations regarding experimental temperature, are provided in the Supplementary Materials (S1).

**Fig 2.**
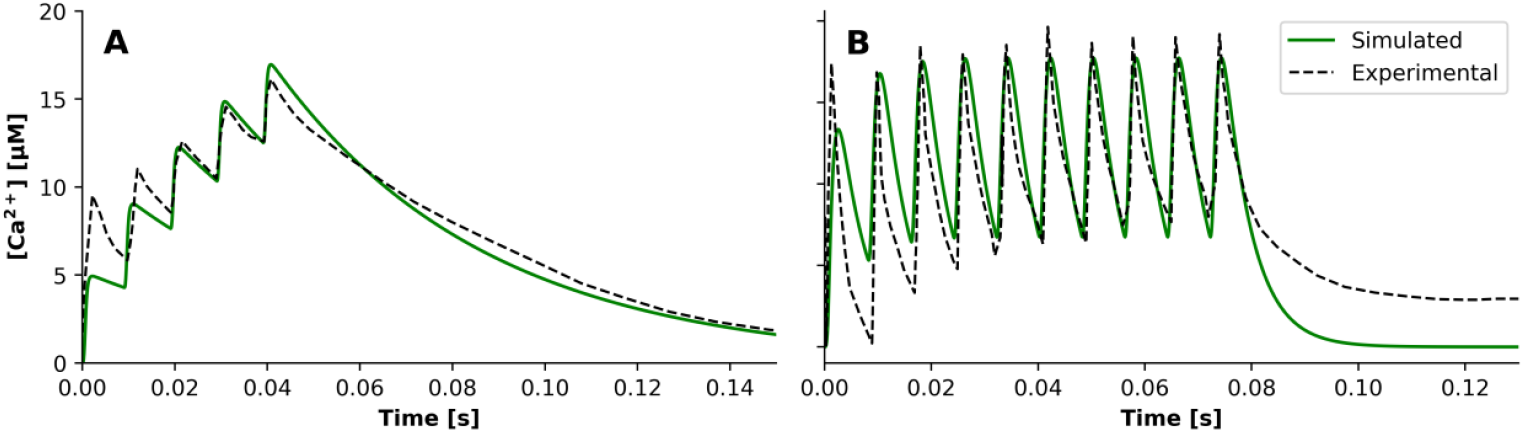
Comparison of the simulated and digitised experimental [Ca^2+^] time course in *μM* for (A) slow rodent fibres at 23°C, stimulated at 100 Hz at constant sarcomere length (corresponding to the Contractile Element length *l*^*CE*^) = 3.8*μm* ≈ 1.6 times the optimal sarcomere length 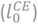 [30, 31] and (B) fast rodent fibres at 35°, stimulated at 125 Hz at 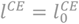. The simulated traces were obtained using the excitation dynamics of [18] and Eq. 1 with binary input trains at the corresponding stimulation frequency.

Unlike [18], the calcium-troponin binding dynamics in the myoplasm were not modelled explicitly (see Supplementary Materials S1). The [Ca^2+^] concentration *c*(*t*) was directly linked to a(t) through a 1^st^ order ODE (Eq. 2) adapted from [12], where the deactivation phase was modified to allow faster decay rates as observed in experimental data:

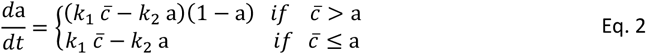

In Eq. 2, 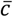 is the calcium concentration normalised by its maximum *c*_*max*_, while *k*_1_and *k*_2_ are the rates of calcium binding to and release from the contractile filament calcium-binding sites. *c*_*max*_, *k*_1_and *k*_2_ assume different values when modelling slow and fast fibres, to account for fibre-type differences in calcium binding capacity [33], and when modelling a muscle and a MU, to reflect the lumping of heterogeneous MU properties into a single actuator at the muscle scale [18]. Importantly, *c*_*max*_, *k*_1_and *k*_2_ are the only parameters that differ between MU- and muscle-scale implementations and the only parameters within the excitation-activation dynamics not directly derived from experimental studies and thus requiring calibration.

#### 2.1.2 PE, SE, and contraction dynamics

The model of [18] was also extended to add the effect of passive tissues, FG pennation angle *α*, an elastic tendon, a FV relationship and the effects of sag and yielding. The 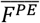, representing the effect of passive connective tissues, was modelled with a 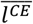-dependent exponential function (Fig 3A), based on [34]. The 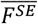 due to an elastic tendon, was modelled as a non-linear function of the tendon strain *ε*^*SE*^ [34] (Fig 3B), additionally including a term to prevent the tendon from going slack [35]. The 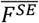 equations include a 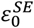 parameter setting the strain at *F*_0_, which was considered species-specific in this work.

**Fig 3.**
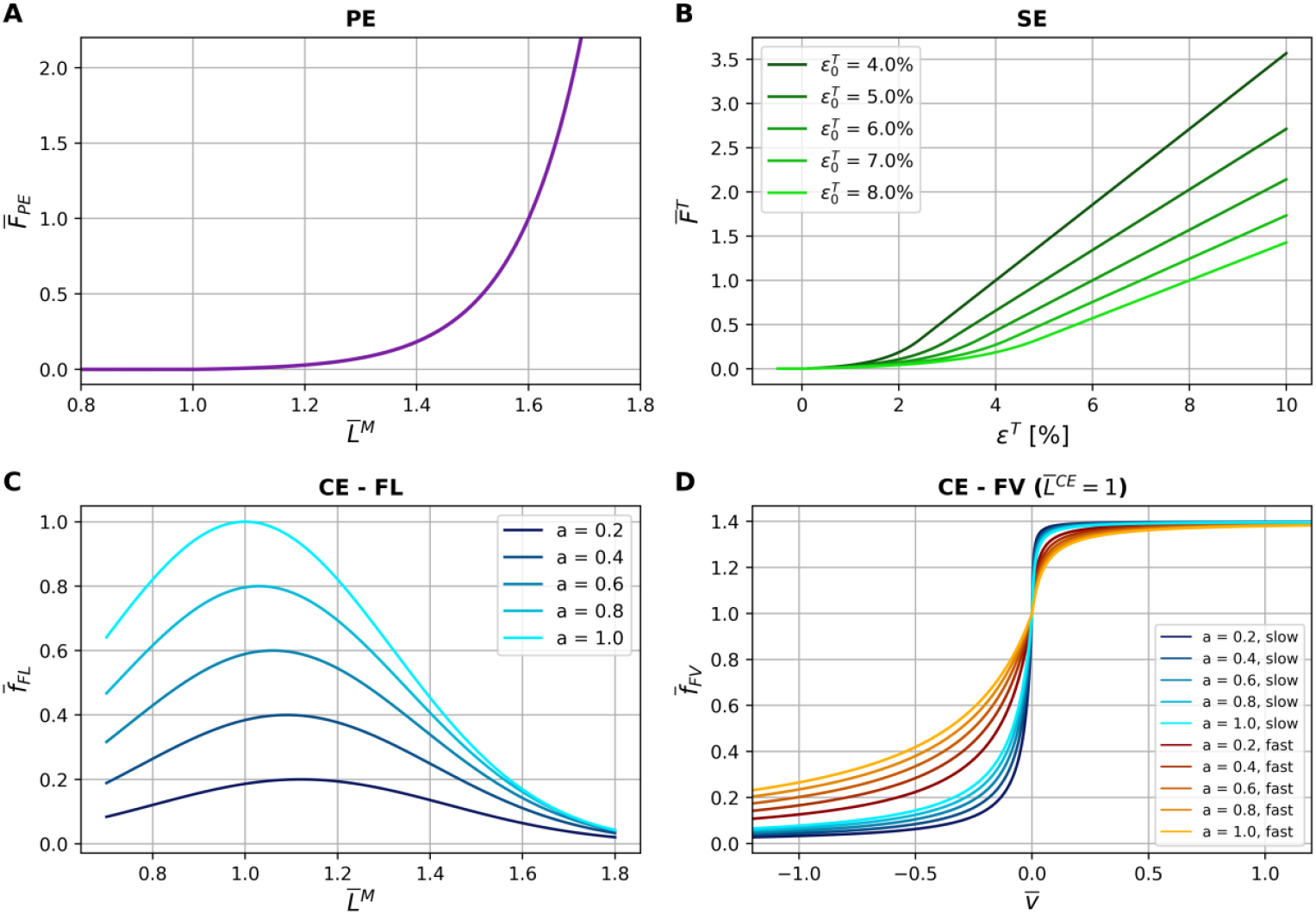
Relationships characterising the model’s (A) Parallel-Elastic Element (PE), (B) Series-Elastic Element (SE) with different values of the species-dependent parameter 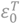, (C) Force-Length (FL) at different activation (a) levels (D) and Force-Velocity (FV) at length 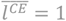 and different a levels.

A FL relationship (Fig 3C), including the activation-dependent shift in optimal fibre length 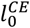 (a), previously reported at sub-maximal activation [36, 37], was also modelled as shown to improve force prediction accuracy [38]. This was used to model the isometric 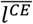-dependent active force component 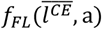 of the CE [39].

To model the dynamic 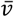-dependent CE’s active force component, a fibre-type-specific FV relationship *f*_*FV*_ (Fig 3D), non-linearly dependent on 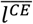 and a(*t*) was adapted from [40] as a variant of Hill’s equation [7] (Eq. 3, Eq. 4).

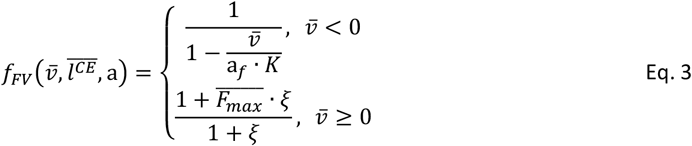

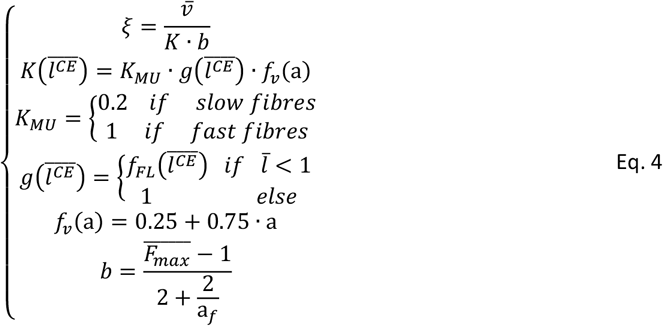

During the concentric phase, the curvature of *f*_*FV*_ is controlled by a_*f*_, which is fibre-type-dependent, as shown experimentally in rodents [41]. In the eccentric phase, the curvature of *f*_*FV*_ is instead controlled by the dimensionless velocity term 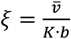, that scales how rapidly force approaches the maximal lengthening force 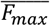. In both phases, *K* represents the maximum contraction velocity under sub-maximal activation. *K* is fibre-type specific through *K*_*MU*_, with fast fibres (IIA fibres in humans) having a value five times higher than slow ones [42]. Consistent with experimental cat data [43], *K* was further reduced at sub-optimal 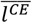 and sub-maximal activation through the scaling factors 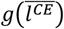 [44] and *f*_*v*_(a) [34], respectively. Finally, the maximum normalised force reachable during lengthening 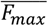 and *v*_*max*_ were set to 1.4 and 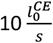 [34], respectively.

#### 2.1.3 Yielding and sag

To better reproduce fibre-type-specific behaviour under experimental conditions, at both MU and muscle scales, yielding [45, 46] (for slow fibres) and sag [4, 47] (for fast fibres), were incorporated into the model. Among others fibre-type-specific effects described in the literature (e.g., susceptibility to fatigue [4] and post-tetanic potentiation [5]), yielding and sag were chosen here because they are present in the contraction protocols simulated in this study (i.e., unfused isometric tetani and dynamic length changes; see Paragraph 2.2), and they are defined as follows. Yielding is a transient reduction in force generation that occurs during rapid changes in muscle length (both lengthening and shortening) at sub-maximal *f* ^*s*^, most prominently in slow fibres. It was modelled as a 1^st^-order ODE with state variable *Y*(*t*) (Eq. 5).

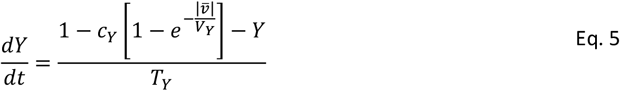

with *c*_*Y*_ = 0.35, *V*_*Y*_ = 0.1 and *T*_*Y*_ = 0.2 [45]. In [45], *Y* scaled the input *f* ^*s*^ in the activation-frequency relationship. Here, as *f* ^*s*^ is experimentally imposed, *Y* was applied instead as a multiplicative factor *φ* to the active state of slow fibres for stimulation frequencies lower than the maximal tetanic frequency 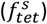 (Eq. 8, Eq. 9). This formulation effectively reduces the number of force-generating cross-bridges during length changes while keeping the experimental input unchanged.

Sag is a characteristic of fast fibres that features a slow force decline during isometric unfused tetanic contractions at low stimulation frequencies [4]. It was modelled as a first-order ODE with state variable *S*(*t*) (Eq. 6, Eq. 7).

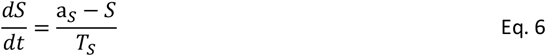

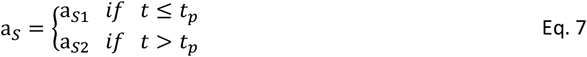

where *T*_*S*_ is a phenomenological time constant, while a_*S*1_ and a_*S*2_ (ranging between 0 and 2) scale the active-state modulation before and after the onset of sag. The onset time *t*_*p*_ was the last local force peak that remained higher than the final force level at the end of stimulation, i.e. the point after which force began its sustained decline. If no such maximum was present, sag was not included. This formulation avoids the arbitrary non-zero threshold of [47]. As for yielding, the sag factor *S* was applied as a multiplicative factor *φ* to the active state, rather than to *f* ^*s*^ as in [47], of fast fibres for *f* ^*s*^ lower than 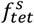 (Eq. 8, Eq. 9).

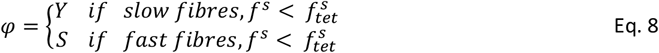

#### 2.1.4 Solving the muscle-tendon dynamics

Assuming tendon elasticity and negligible muscle inertial forces, the muscle-tendon dynamics are expressed by the following equation (Eq. 9).

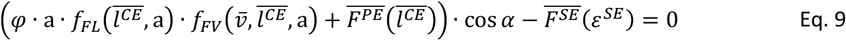

To calculate *F*^*CE*^ in a forward dynamic simulation, 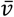 is isolated as an explicit state equation (Eq. 10) and integrated in time to update 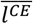.

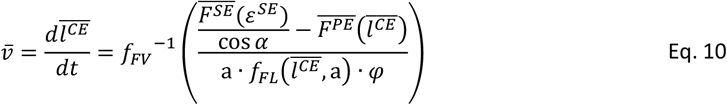

where *f*_*FV*_ ^−1^ is the inverse of Eq. 10. To avoid singularities, activation was kept above a small positive threshold (a(*t*) ≥ a_*min*_), *f*_*FV*_ (*t*) constrained to its admissible range 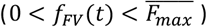 and the pennation was restricted such that cos *α*(*t*) > 0, thereby avoiding singularities associated with pennation angles approaching 90°.

The system of ODEs consisting of the excitation (see Supplementary Materials S1), activation (Eq. 1, Eq. 2) and contraction dynamics (Eq. 3Eq. 7), with the state variables *u, c* and their first derivative, a,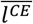 and *φ* (when included), was solved at each time step as an initial value problem. The proposed model, structurally summarised in Fig 4, was implemented in Python, using the LSODA method [48] from the SciPy 1.0 library [49] to integrate the system of ODE.

**Fig 4.**
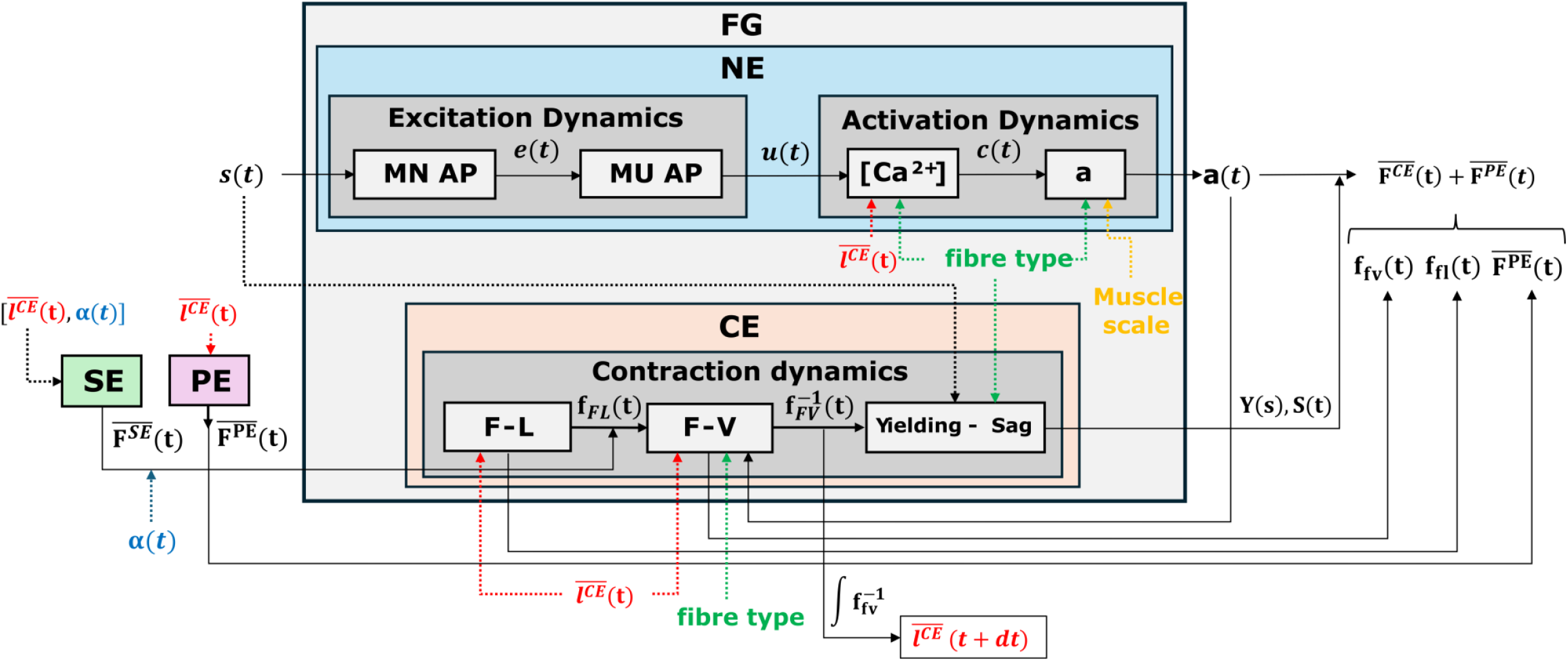
Comprehensive diagram of the structural connections and variable flow of the proposed Hill-type model. The Force Generator (FG) comprises a Neuromechanical Element (NE) and a Contractile Element (CE). The NE includes the excitation-activation dynamics, receiving the neural input *s*(*t*) and producing the active state a(*t*). The intermediate stage modelling the [*Ca*^2+^] dynamics depends on the CE length 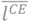 and on the fibre type, which also affects the dynamics of a(*t*). The CE includes the Force-Length (FL) and Force-Velocity (FV) relationships (producing forces *f*_*FL*_(*t*) and *f*_*FV*_ (*t*)), both dependent on 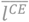, with FV additionally dependent on fibre type and a(*t*). The CE also includes the time-dependent effects of yielding *Y*(*t*) and Sag *S*(*t*), which act multiplicatively on a(*t*). The normalised active force of the CE, 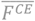, is thus computed as the product of the modulated a(*t*), *f*_*FL*_(*t*), and *f*_*FV*_ (*t*), while the integration of 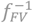 determine 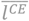 at the next time step *t* + *dt*. Externally to the FG, the Parallel-Elastic Element (PE) and Series-Elastic Element (SE) produce passive forces 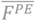 and 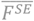, respectively, both dependent on 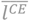.

### 2.2 Model benchmarks

Model performance was evaluated against four benchmark datasets derived from both previously published studies and ad hoc experiments. These datasets were designed to span the two actuator scales (MU and whole muscle), the two main contractile phenotypes (i.e., slow- and fast-twitch MUs; slow and fast muscles), and both isometric and dynamic conditions where available. Specifically, the benchmarks comprised datasets spanning slow and fast MUs from cat and rat lateral (LG) and medial (MG) gastrocnemius under isometric conditions, slow muscles from cat and rat soleus under isometric and dynamic conditions and fast muscles from rat extensor digitorum longus (EDL) and cat caudofemoralis (CF) under isometric and dynamic conditions, respectively.

Benchmarks were identified by *B* with superscript *MU* or *M* indicating the actuator scale (MU or muscle), suffix −*S* or −*F* indicating slow or fast fibre composition and subscript *iso* or *dyn* indicating isometric and dynamic contractions. For isometric benchmarks (*B*_*iso*_), the additional specifier −*f* indicates constant length trials at varying *f* ^*s*^ (force-frequency validation), while −*l* indicates constant-*f* ^*s*^ trials at different fixed lengths (force-length validation).

#### 2.2.1 MU benchmarks

The MU benchmarks 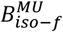 trials were based on *in vivo* quasi-isometric single-MU experiments conducted on the cat [4] and rat [50] MG and LG. In these studies, MUs were categorised as Slow, Fast Fatigue-Resistant and Fast-Fatigable (FF). Here, a single fast MU category was considered, parameterised from Fatigue-Resistant units, as they are more representative of the continuous operating spectrum of mammalian skeletal muscles [3]. In total, eleven force traces were manually digitised from [4, 50] to constitute the 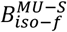 and 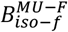 benchmarks (Table 1). The 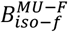 benchmarks were further subdivided into 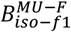 and 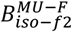 to distinguish between the cat and rat MG, respectively.

**Table 1.**
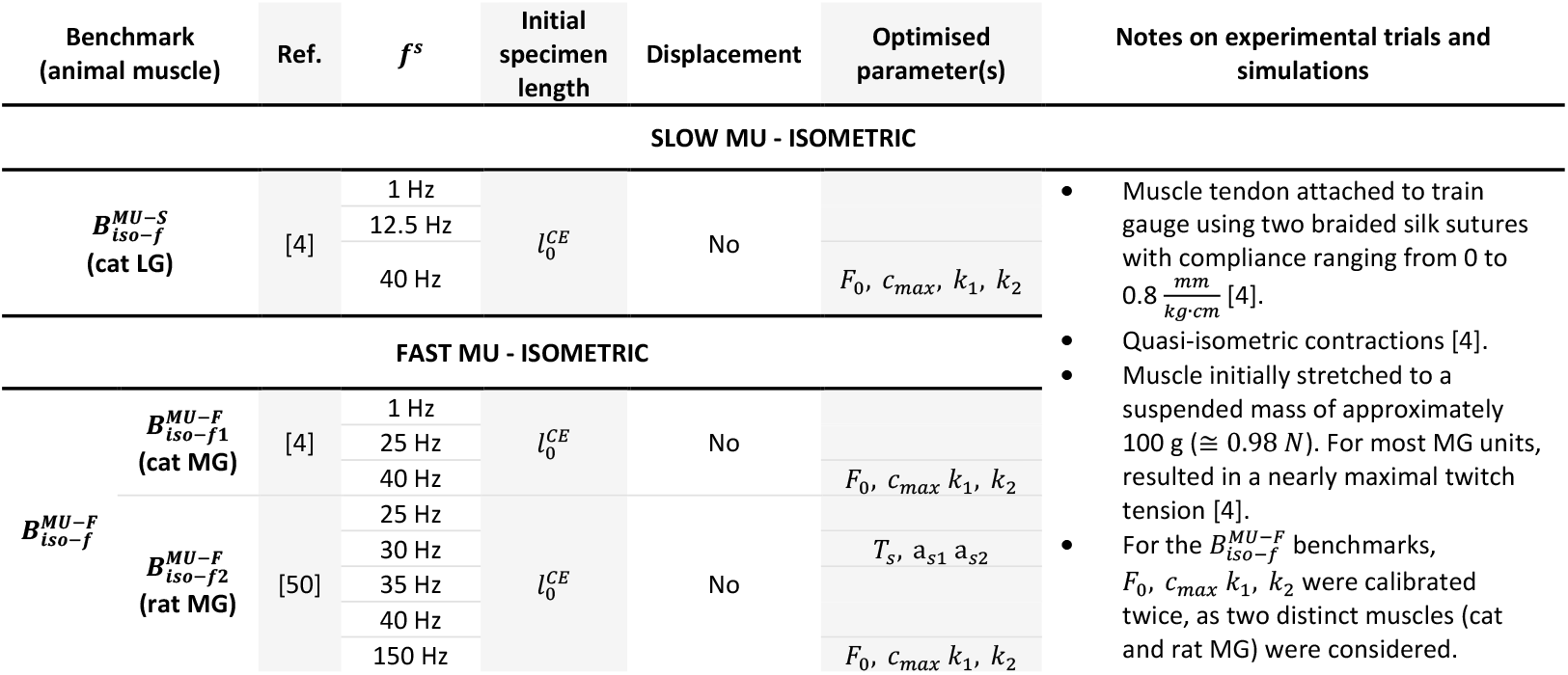
Experimental trials included in the isometric slow 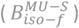 and fast 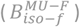 motor-unit scale benchmarks at varying stimulation frequencies. This table summarises the *in vivo* validation trials performed on single motor units (MUs) of the cat [4] and rat [50] lateral gastrocnemius (LG) and medial gastrocnemius (MG). All trials were conducted under isometric conditions (no displacement) at length 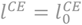. The stimulation frequency (*f* ^*s*^) values ranged from 1 Hz for single twitches up to 40 Hz for maximum tetanic force. The second column from the right highlights the specific model parameters calibrated via optimisation against each respective dataset.

To simulate these experimental conditions (Table 1), the tendon was considered rigid to represent stiff tendon-strain gauge sutures [4] and the complete tendon excision [50]. The FV relationship was neglected because of the isometric contractile conditions and length was fixed at 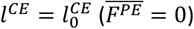 because of the isometric pre-loaded preparation. Under these assumptions, Eq. 9 reduces to Eq. 11, with Eq. 8 unchanged.

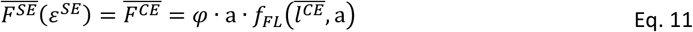

No dynamic MU benchmark was included because, to the authors’ best knowledge, the available studies that measured the FV relation of individual MUs [51-53] did not report enough traces capable of being digitised to constitute a complete benchmark.

#### 2.2.2 Slow muscle benchmarks

The slow muscle benchmarks *B*^*M*−*S*^ (Table 2) were based on *in vivo* experiments conducted on the cat [14] and rat [23] soleus muscle-tendon unit.

**Table 2.**
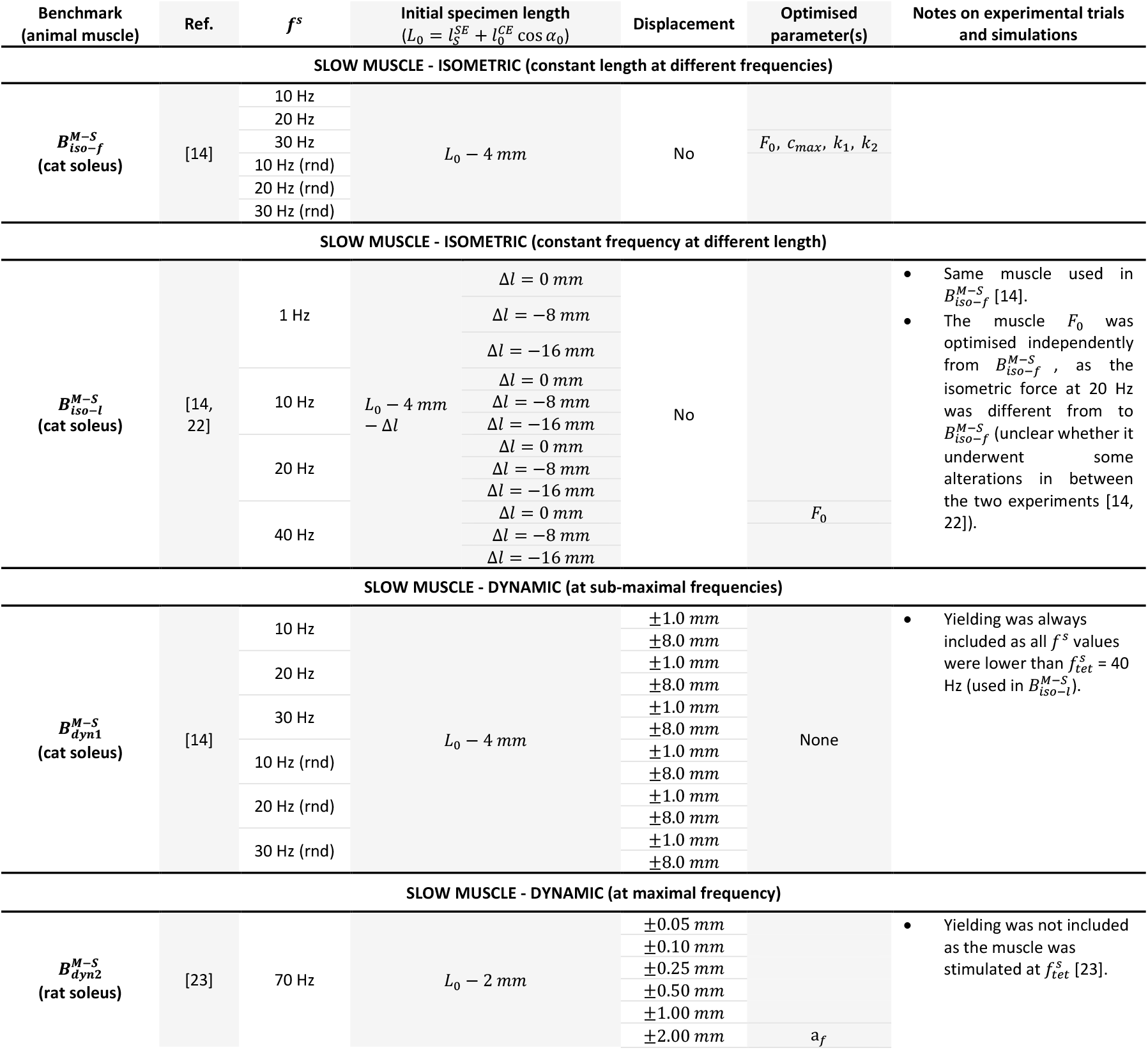
Experimental trials included in the slow whole-muscle scale benchmarks (*B*^*M*−*S*^). This table summarises the *in vivo* validation trials performed on slow cat [14] and rat [23] soleus muscles. The benchmarks are categorised into constant and randomised stimulation frequency (*f* ^*s*^) isometric tests 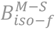 [14], length-varying isometric tests 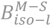 [14] and dynamic tests 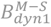 [14, 23] and 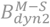 [23]. The second column from the right highlights the specific model parameters calibrated via optimisation against each respective dataset.

The cat soleus, initially deactivated, was stimulated at sub-maximal constant *f* ^*s*^ (10, 20 and 30 Hz) and random *f* ^*s*^ patterns with the same mean frequencies under isometric conditions 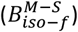 [15] and during imposed muscle-tendon length changes of ±1 and ±8 mm 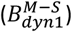. The same muscle was also stimulated at constant *f* ^*s*^ (1, 10, 20 and 40 Hz) at three different lengths 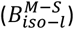, with force traces reported in [22]. To model the cat soleus it was assumed 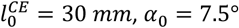 and 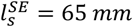, as measured by [54], and 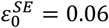 was used to represent the high compliance of the cat soleus tendon described in [55].

The rat soleus benchmark 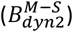 consisted of six dynamic trials from [15] in which the muscle, initially deactivated, was maximally stimulated at 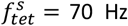 and subjected to controlled progressive stretches increasing from 0.05 to 2.0 mm (Table 2), representative of physiological locomotor movements. To model the rat soleus it was assumed 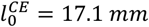 as measured by [23], while 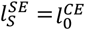 and *α*_0_ = 6° from [56]. The same 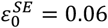 used for the cat soleus was adopted, consistent with the peak rat soleus tendon strains reported in [57].

#### 2.2.3 Fast muscle benchmarks

The fast muscle benchmarks *B*^*M*−*F*^ (Table 3) comprised isometric data from a single rat extensor digitorum longus (EDL) muscle, collected ad hoc for this study at the University of Stuttgart, and dynamic data from the cat caudofemoralis (CF) reported in [58]. The EDL experiments were conducted in accordance with Section 8 of the German Animal Welfare Act (Tierschutzgesetz, §4(3); permit no. T 236/24; additional details of the in vitro experimental protocol are provided in Supplementary Materials S4).

**Table 3.**
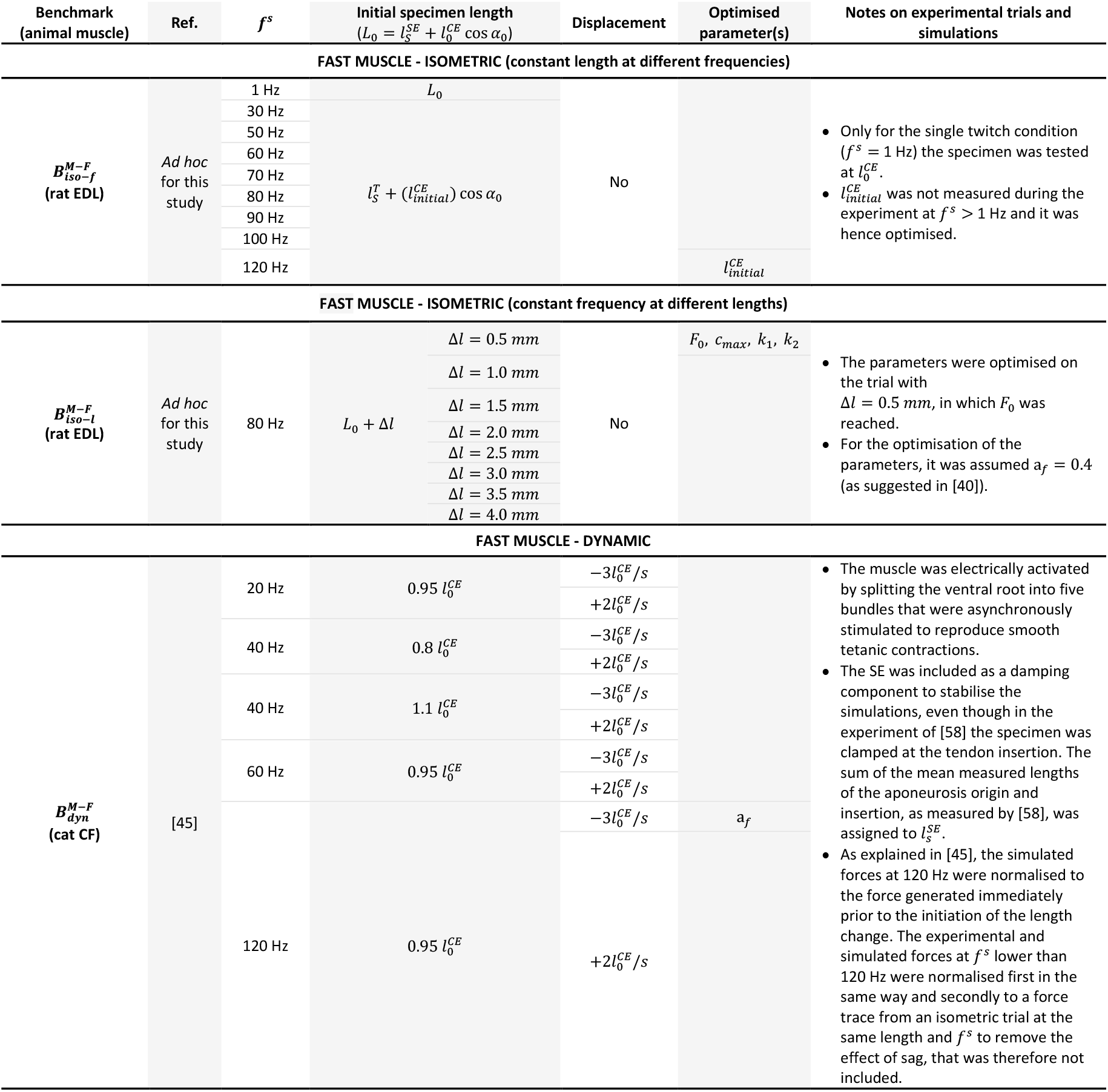
Experimental trials included in the fast whole-muscle scale benchmarks (*B*^*F*−*S*^). This table details the experimental conditions used to validate the model against fast rat extensor digitorum longus (EDL) *in vitro* (*ad hoc* for this study) and cat caudofemoralis (CF) *in vivo* [45]. The benchmarks are categorised into constant stimulation frequency (*f* ^*s*^) isometric tests 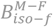, length-varying isometric tests 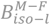 and dynamic tests 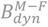 [45]. The second column from the right highlights the specific model parameters calibrated via optimisation against each respective dataset.

For the isometric benchmark on the rat EDL, initial isometric normalisation trials were conducted to determine 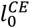 (6.66 mm) and *F*_0_ (2.49 N). The muscle was then tested isometrically at a constant muscle-tendon unit length under nine increasing *f* ^*s*^ values from 1 to 120 Hz (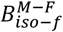, Table 3) and at eight different fixed lengths under constant *f* ^*s*^ at 80 Hz 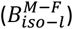. To model the rat EDL, 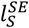 was set equal to the distal tendon left for testing (5.0 mm), *α*_0_ = 10° based on previous rat EDL measurements [59] and 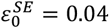 consistently with published rat EDL series-elasticity data [60] and the original value in [34].

The dynamic fast muscle benchmark 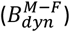 was based on *vivo* experiments on the cat CF [58]. In these experiments, the muscle was electrically activated at different *f* ^*s*^ values and subjected to shortening and lengthening from different initial specimen lengths 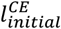. Five experimental conditions, manually digitised from [45] yielded ten force traces in total (shortening and lengthening for each condition; see Table 3). For the cat CF, the model parameters were set to values measured or derived from [58]: *F*_0_ = 15.4 N, 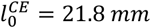 and *α*_0_ = 0° (measured), 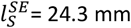 (derived, see notes in Table 3) and 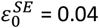 (derived, consistent with the little in-series elasticity description).

#### 2.2.4 Parameters calibration

All model parameters not explicitly calibrated below were fixed from previous studies or from the benchmark-specific experimental conditions, as described in the previous sections and in Supplementary Materials S1-S3. In each benchmark, the remaining unknown parameters were calibrated using a Nelder-Mead optimisation algorithm [61], with an objective function that minimised the sum of squared errors between the simulated force output of our model and the experimental force from the benchmark dataset.

The active-state parameters *c*_*max*_, *k*_1_and *k*_2_ (Eq. 2), together with *F*_0_ when required, were calibrated for each modelled muscle species from the isometric trial at the maximum available constant *f* ^*s*^. This trial is representative of maximal force-generating capacity under nearly saturated [Ca^2+^] conditions (i.e. *c* ≈ *c*_*max*_), in which the force profile is primarily determined by the activation dynamics with minimal influence of the FL and FV relationships.

The FV relationship parameter a_*f*_ (Eq. 3, Eq. 4) was calibrated from the trial combining the maximum *f* ^*s*^ and the largest displacement amplitude, where velocity-dependent effects were expected to be maximal at near-constant activation. Because the dependence of sag on *f* ^*s*^ is not fully established, the sag parameters *T*_*s*_, a_*s*1_ and a_*s*2_ (Eq. 6, Eq. 7) were calibrated from the isometric trial at a stimulation frequency closest to the mean of the submaximal *f* ^*s*^ values. The complete set of optimised parameters is reported in Supplementary Materials S3 (Table S1).

#### 2.2.5 Model evaluation

Model predictive performance was first evaluated for all benchmarks by comparing simulated and experimental force traces (including the ones used for calibrating the model parameters) using the mean absolute error (mAE), maximum absolute error (MAE) and coefficient of determination (*R*^2^). Errors were expressed as percentage of *F*_0_ and, unless otherwise stated, were calculated over the full duration of each force trace. For the dynamic fast-muscle benchmark 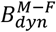, evaluation was restricted to the interval of imposed length change.

Additional condition-specific metrics were then computed for the relevant isometric trials. For all isometric trials at constant *f* ^*s*^, peak force *F*^*pk*^ was also computed over the full trial duration [50] and the corresponding peak-force error Δ*F*^*pk*^ was computed as the difference between simulated and experimental *F*^*pk*^, expressed as a percentage of *F*_0_. For isometric twitch and fused-tetanus trials at constant *f* ^*s*^, the temporal descriptors *τ*_*rise*_ and *τ*_*decay*_ were additionally evaluated. For twitch contractions, *τ*_*rise*_ was defined as the time from force onset to *F*^*pk*^, and *τ*_*decay*_ as the time from *F*^*pk*^ to 50% force decay [50]. In fused tetani, equivalent first-order time constants were used, corresponding to the time required to reach 63.2% (*τ*_*rise*_) and decay to 36.8% (*τ*_*decay*_) of *F*^*pk*^ [8]. For isometric unfused tetanus trials at constant *f* ^*s*^, the fusion index (FI) was defined as the ratio between the minimum force between the last two consecutive force peaks and the last peak force [62]. The errors Δ*τ*_*rise*_, Δ*τ*_*decay*_ and ΔFI were then computed as the difference between the simulated and experimental *τ*_*rise*_, *τ*_*decay*_ and FI.

For benchmarks in which yielding or sag influenced force production, simulations were repeated with and without these mechanisms to quantify their contribution to the model accuracy. Their effect on model accuracy was assessed using a paired non-parametric Wilcoxon signed-rank test across all error metrics.

## 3 Results

Table 4 and Table 5 summarise the model performance across all benchmarks. Table 4 reports the global waveform-fit metrics (mAE, MAE and *R*^2^) for all simulated conditions. Table 5 reports the additional metrics (Δ*τ*_*rise*_, Δ*τ*_*decay*_, ΔFI and Δ*F*^*pk*^) for isometric conditions. Across all simulations, the model reproduced the main features of force production with mean errors generally below 15% *F*_0_, although higher errors (up to ∼20-25%) were observed in specific conditions. Model accuracy was consistently higher at moderate-to-high stimulation frequencies (≥ 20 Hz), while larger discrepancies were observed at low frequencies, particularly under twitch and unfused-tetanus conditions. The inclusion of fibre-type-specific mechanisms had distinct effects depending on the benchmark: yielding significantly improved the prediction of dynamic force responses in slow muscles, whereas sag had a more variable influence on the accuracy of fast MU simulations, with no consistent improvement across conditions. An extended version of both Table 4 and Table 5 with the results of individual benchmark trials can be found in Supplementary Materials S5-S7.

**Table 4.** Quantitative results summarising the model’s predictive accuracy. This table provides a comprehensive evaluation of the model’s performance by comparing simulated forces to experimental measurements across all tested conditions. Accuracy is expressed through the mean ± standard deviation, minimum, and maximum values of the mean absolute error (mAE), maximum absolute error (MAE) and coefficient of determination (*R*^2^). For mAE and MAE, results are expressed as % of the maximum isometric force *F*_0_. The table additionally reports the change in predictive error obtained by incorporating yielding (Y) in the model, with statistically significant changes indicated by an asterisk (*) based on the Wilcoxon signed-rank test.

| Benchmark<br>(n. trials) | mAE Mean ± SD<br>[% $F_0$] | mAE Min. - Max.<br>[% $F_0$] | MAE Mean ± SD<br>[% $F_0$] | MAE Min. - Max.<br>[% $F_0$] | $R^2$ Mean ± SD | $R^2$ Min. - Max. |
| --- | --- | --- | --- | --- | --- | --- |
| **SLOW MU** |  |  |  |  |  |  |
| $B_{iso-f}^{MU-S}$ (3) | 3.8 ± 3.9 | 0.8 - 9.3 | 13.4 ± 7.7 | 5.2 - 23.8 | 0.8 ± 0.1 | 0.6 - 0.9 |
| **FAST MU** |  |  |  |  |  |  |
| $B_{iso-f}^{MU-F}$ (8) | 6.7 ± 6.0 (no S)<br>6.7 ± 5.9 (S) | 1.6 - 19.1 (no S)<br>1.6 - 18.5 (S) | 25.6 ± 9.2 (no S)<br>24.5 ± 8.7 (S) | 16 - 43.5 (no S)<br>16 - 41.3 (S) | 0.7 ± 0.2 (no S)<br>0.7 ± 0.2 (S) | 0.1 - 0.9 (no S)<br>0.2 - 0.8 (S) |
| **SLOW MUSCLE** |  |  |  |  |  |  |
| $B_{iso-f}^{M-S}$ (6) | 5.0 ± 3.4 | 0.96 - 10.3 | 13.0 ± 5.6 | 5.5 - 19.9 | 0.0 ± 2.0 | (-4.0) - 0.9 |
| $B_{iso-l}^{M-S}$ (12) | 8.3 ± 8.1 | 0.2 - 24.7 | 20.4 ± 13.3 | 1.2 - 43.9 | 0.5 ± 0.4 | (-0.4) - 0.9 |
| $B_{dyn1}^{M-S}$ (12) | 10.8 ± 3.8 (no Y)<br>7.9 ± 2.4 (Y*) | 5.2 - 18.3 (no Y)<br>4.1 - 11.6 (Y) | 37.1 ± 16.3 (no Y)<br>27.2 ± 8.8 (Y*) | 18.6 - 67.1 (no Y)<br>14.8 - 41.4 (Y) | (-1.1) ± 3.4 (no Y)<br>(-0.4) ± 2.5 (Y*) | (-10) - 0.9 (no Y)<br>(-6.4) - 0.9 (Y) |
| $B_{dyn2}^{M-S}$ (6) | 4.9 ± 2.5 | 1.7 - 8.7 | 23.4 ± 11.4 | 10.1 - 42.5 | 0.9 ± 0.0 | 0.8 - 0.9 |
| **FAST MUSCLE** |  |  |  |  |  |  |
| $B_{iso-f}^{M-F}$ (12) | 6.5 ± 3.3 | 1.5 - 10.8 | 19.5 ± 8.1 | 4.5 - 26.3 | 0.8 ± 0.1 | 0.6 - 0.9 |
| $B_{iso-l}^{M-F}$ (8) | 5.0 ± 1.1 | 3.5 - 6.7 | 16.4 ± 6.1 | 11.1 - 30.4 | 0.9 ± 0.0 | 0.7 - 0.9 |
| $B_{dyn}^{M-F}$ (10) | 15.0 ± 6.4 | 2.4 - 24.1 | 31.2 ± 11.7 | 7.0 - 46.8 | (-0.6) ± 1.4 | (-4.4) - 0.9 |

**Table 5.**
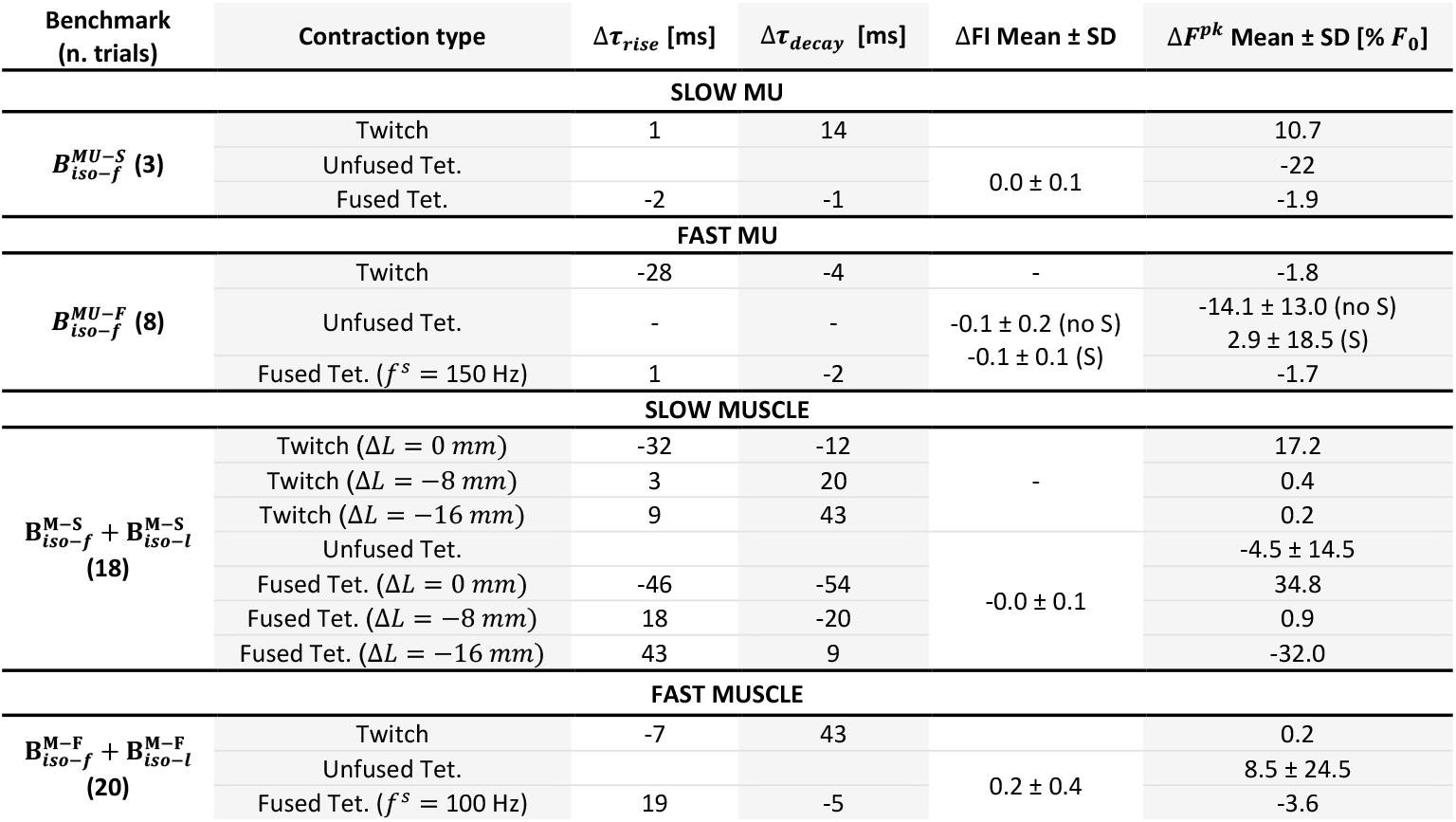
Quantitative results summarising the model’s predictive accuracy comparing simulated forces to experimental measurements across all isometric conditions. For twitch and fused-tetanus contractions, accuracy was assessed from the differences between simulated and experimental rise and decay time constants (Δ*τ*_*rise*_ and Δ*τ*_*decay*_). For all conditions, the difference between simulated and experimental peak force (Δ*F*^*pk*^) was computed as percentage of the maximum isometric force *F*_0_ and reported as mean ± standard deviation among trials (unless a single trial is considered). For unfused-tetanus conditions only, differences between the simulated and experimental fusion index (ΔFI) were also calculated and reported as mean ± standard deviation among trials (unless a single trial is considered). The table additionally reports the change in predictive error obtained by incorporating sag (S) in the model, with statistically significant changes indicated by an asterisk (*) based on the Wilcoxon signed-rank test.

### 3.1 MU benchmarks

The 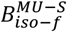 and 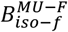 benchmarks were used to assess the model under isometric MU conditions across slow and fast units. Overall, the model reproduced the shape and amplitude of most experimental isometric MU force traces with good accuracy across the considered *f* ^*s*^ values (Table 4, Fig 5). Better performance was obtained for slow than fast MUs: 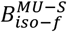 showed mAE = 3.8 ± 3.9%, MAE = 13.7 ± 7.7%, and *R*^2^ = 0.8 ± 0.1, whereas 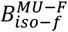 showed mAE = 6.7 ± 6.0%, MAE = 25.6 ± 9.2%, and *R*^2^ = 0.7 ± 0.2 (when sag was not included in the model).

**Fig 5.**
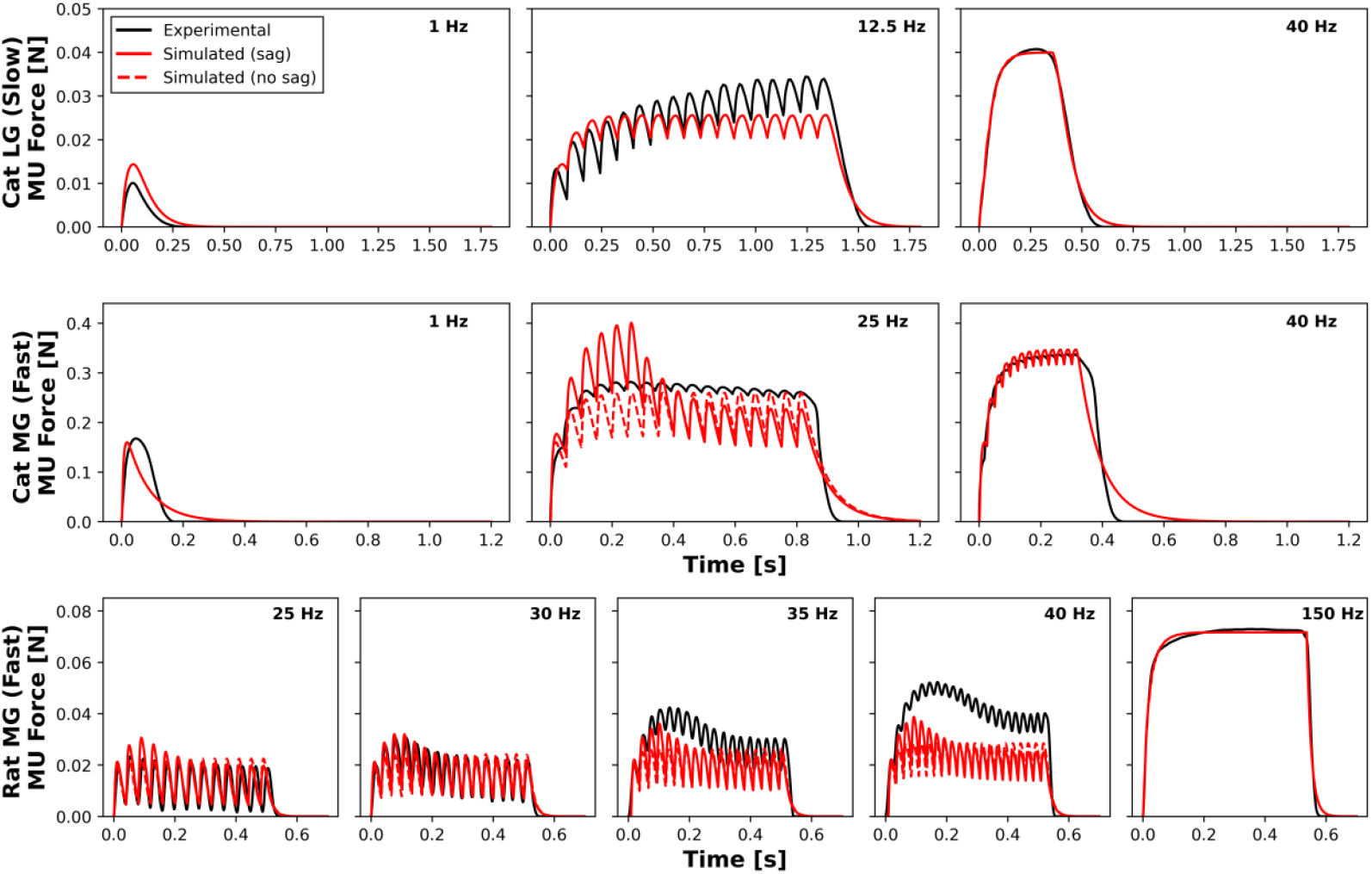
Experimental and simulated isometric force responses of the cat lateral gastrocnemius (LG) slow motor unit (MU) [4] (first row, 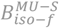 benchmarks), cat medial gastrocnemius (MG) fast MU [4] (second row, 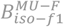 benchmarks) and rat MG fast MU [50] (third row, 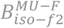 benchmarks) held at optimal fibre length and stimulated at different frequencies. The experimental traces were manually digitised from [4, 50].

Time- and *F*^*pk*^-related errors were largest for twitch and unfused-tetanus conditions (Table 5). For slow MUs, the simulated twitch showed a longer decay phase (Δ*τ*_*decay*_: 14 ms) and summed to an unfused tetanus with a 22% lower *F*^*pk*^ than the experimental trace (Table 5). For fast MUs, the largest errors occurred for the rat GM MU stimulated at 40 Hz, where *F*^*pk*^ was underestimated by 18% and 33% with and without sag, respectively (Fig 5, Table S3 in Supplementary Materials S5), and the simulated twitch peaked earlier than the experimental one (Δ*τ*_*raise*_: 28 ms). Across all MU conditions, FI was generally well reproduced, with the worst-case prediction corresponding to a contraction that was 30% less fused than the experimental one (Fig 5, Table S3 in Supplementary Materials S5). The inclusion of sag in modelling fast MUs did not result in a statistically significant improvement of any error metric (p > 0.06).

### 3.2 Slow muscle benchmarks

#### 3.2.1 Isometric slow muscle benchmarks

To validate the force-frequency relationship of the model’s NE and CE at the slow-type muscle scale, simulations were compared against the 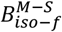 benchmarks (Fig 6A). For the constant-*f* ^*s*^ trials (Fig 6A left column) the errors decreased from 10.3% at 1 Hz, to 3.5% at 20 Hz with increasing *f* ^*s*^ (Fig 6B left column), up to 30 Hz (optimised trial). This error trend was reflected in the random-*f* ^*s*^ trials (Fig 6A right column), with mAE values ranging from 8.7% to 1.9% at 10 and 30 Hz mean *f* ^*s*^ (Fig 6B right column), respectively. The MAEs across all isometric conditions were between 5.2 and 23.8% (Table 4). Notably, the model reproduced the tetanic fusion level with a high accuracy for all trials, with a maximum FI difference of 0.1 (Table S5 in Supplementary Materials S6).

**Fig 6.**
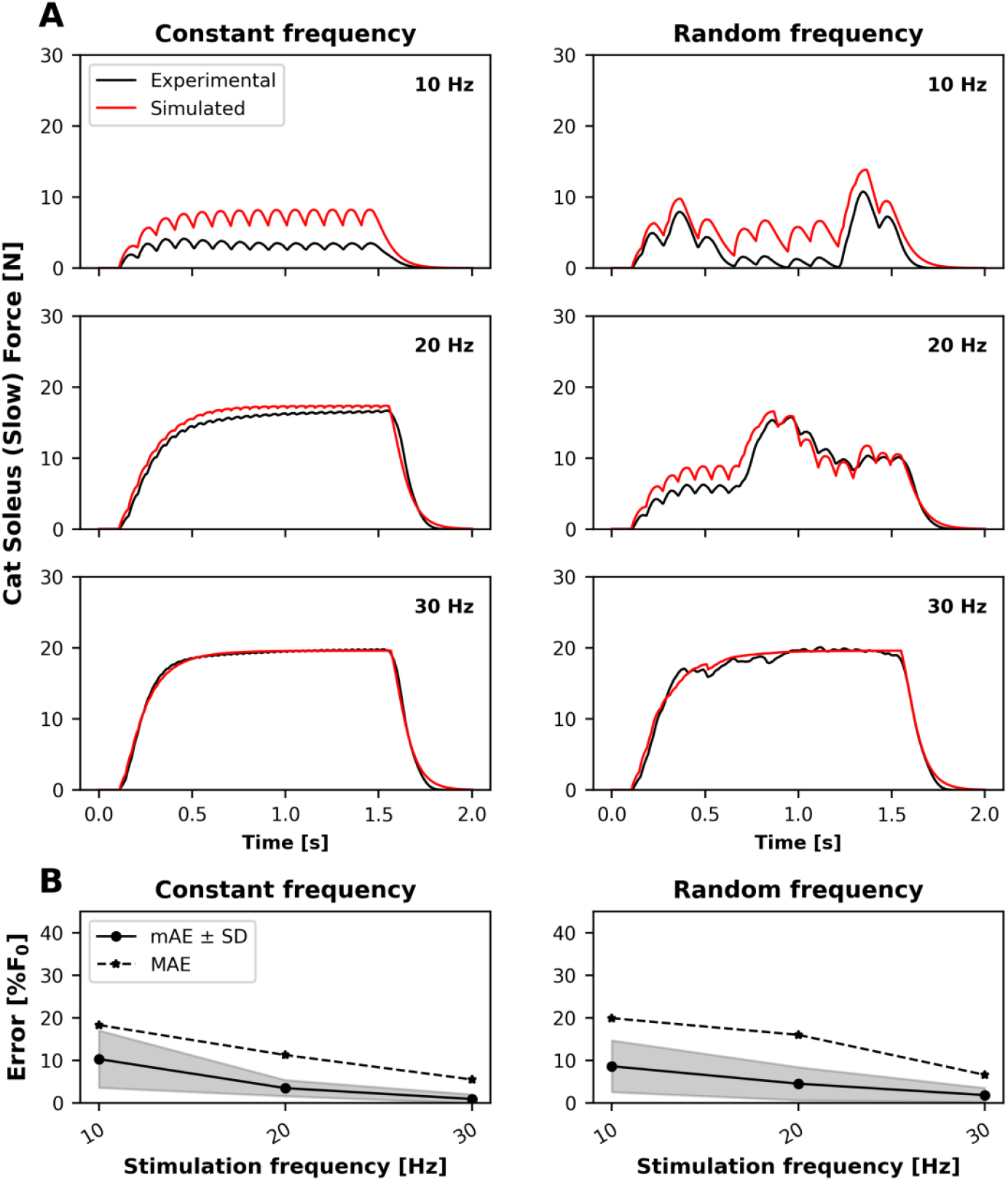
(A) Experimental and simulated isometric force responses of the cat soleus muscle-tendon unit (slow whole-muscle scale; 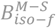 benchmarks) to constant (left column) and random (right column) stimulation patterns with mean stimulation frequencies of 10 Hz (first row), 20 Hz (second row), and 30 Hz (third row), while held at a length of 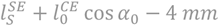 [14]. (B) Mean absolute error (mAE, black solid line) ± standard deviation (SD, grey area) and maximum absolute error (MAE, black dashed line) for the 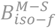 benchmarks.

The 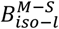 benchmarks were used to validate the FL relationship of the model at the same whole-muscle scale (Fig 7A). The model accurately reproduced the rat soleus experimental twitch responses (1 Hz), with mAE ≤ 2%, at all lengths except at Δ*l* = 0 *mm* (Table 4, Fig 7B). At that length, *F*^*pk*^ was overestimated by 17% and occurred 32 ms earlier than in the experimental trace (Table 5). At higher *f* ^*s*^ values, the smallest errors were obtained near the optimal operating range 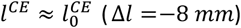, with 0.3% ≤ mAE ≤ 6.9% (Fig 7B central column). At *f* ^*s*^ of 20 Hz and 40 Hz, the predicted *F*^*pk*^ shifted with Δ*l*, from overestimation at Δ*l* = 0 *mm* (up to +35%) to underestimation at Δ*l* = −16 *mm* (up to -25%), as shown in Fig 7A and Fig 7B.

**Fig 7.**
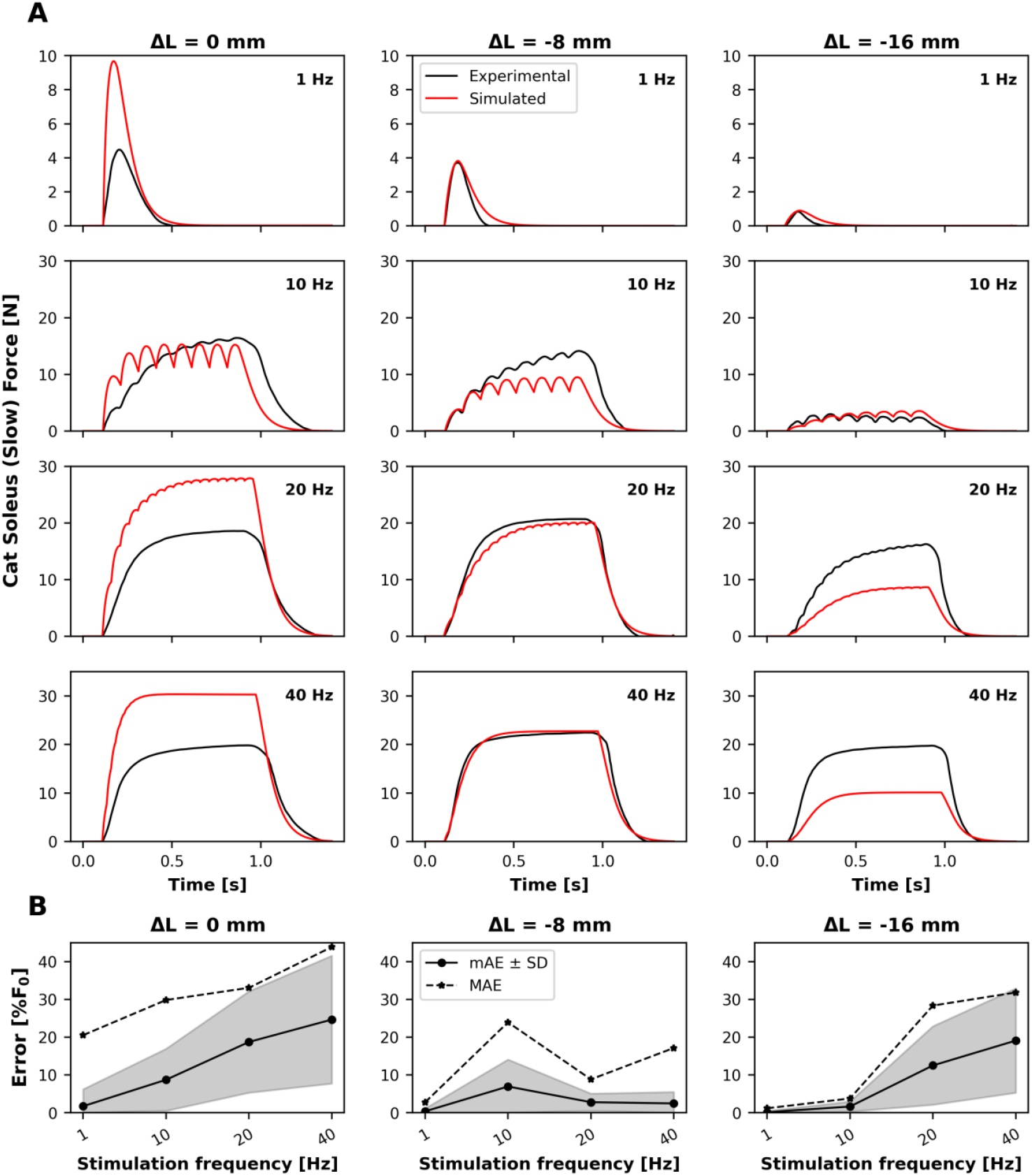
(A) Experimental and simulated isometric twitch (at 1 Hz, first row), unfused tetanic (at 10 and 20 Hz, second and third row) and fused tetanic (at 40 Hz, fourth row) force responses of the cat soleus muscle-tendon unit (slow whole-muscle scale; 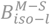 benchmarks) at three lengths defined by Δ*l* = 0, −8, and −16 mm, corresponding to a total muscle-tendon length of 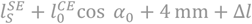. Experimental traces were manually digitised from [22]. (B) Mean absolute error (mAE, solid black line) ± standard deviation (SD, grey shaded area) and maximum absolute error (MAE, dashed black line) for the 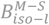 benchmarks.

#### 3.2.2 Dynamic slow muscle benchmarks

The 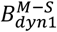 benchmarks (Fig 8A, B) were used to validate the FV relationship of the model under sub-maximal stimulation conditions at the slow-type whole-muscle scale. When yielding was considered, the model reproduced the experimental rat soleus force responses with a mean mAE of 7.9 ± 2.4% and 14.8% ≤ MAE ≤ 41.4% (Table 4), significantly improving model accuracy (p < 0.001) across all stimulation frequencies and displacement amplitudes (Table 4; Fig 8A, B, red solid vs dashed lines). Overall, the largest discrepancies were observed at the lowest stimulation frequency (10 Hz), consistently with what was observed in the 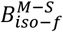 benchmarks. At this frequency, mAE values reached 11.1% for constant *f* ^*s*^ (Fig 8C) and 8.0% for random *f* ^*s*^ (Fig 8D) at ±1 mm displacement amplitude. Increasing the displacement amplitude from ±1 mm (Fig 8A, B, left column) to ±8 mm (Fig 8A, B right column) generally resulted in mAE differences that increased by almost 2%, reaching 5% under random *f* ^*s*^ stimulation at 20 Hz (Fig 8C, D).

**Fig 8.**
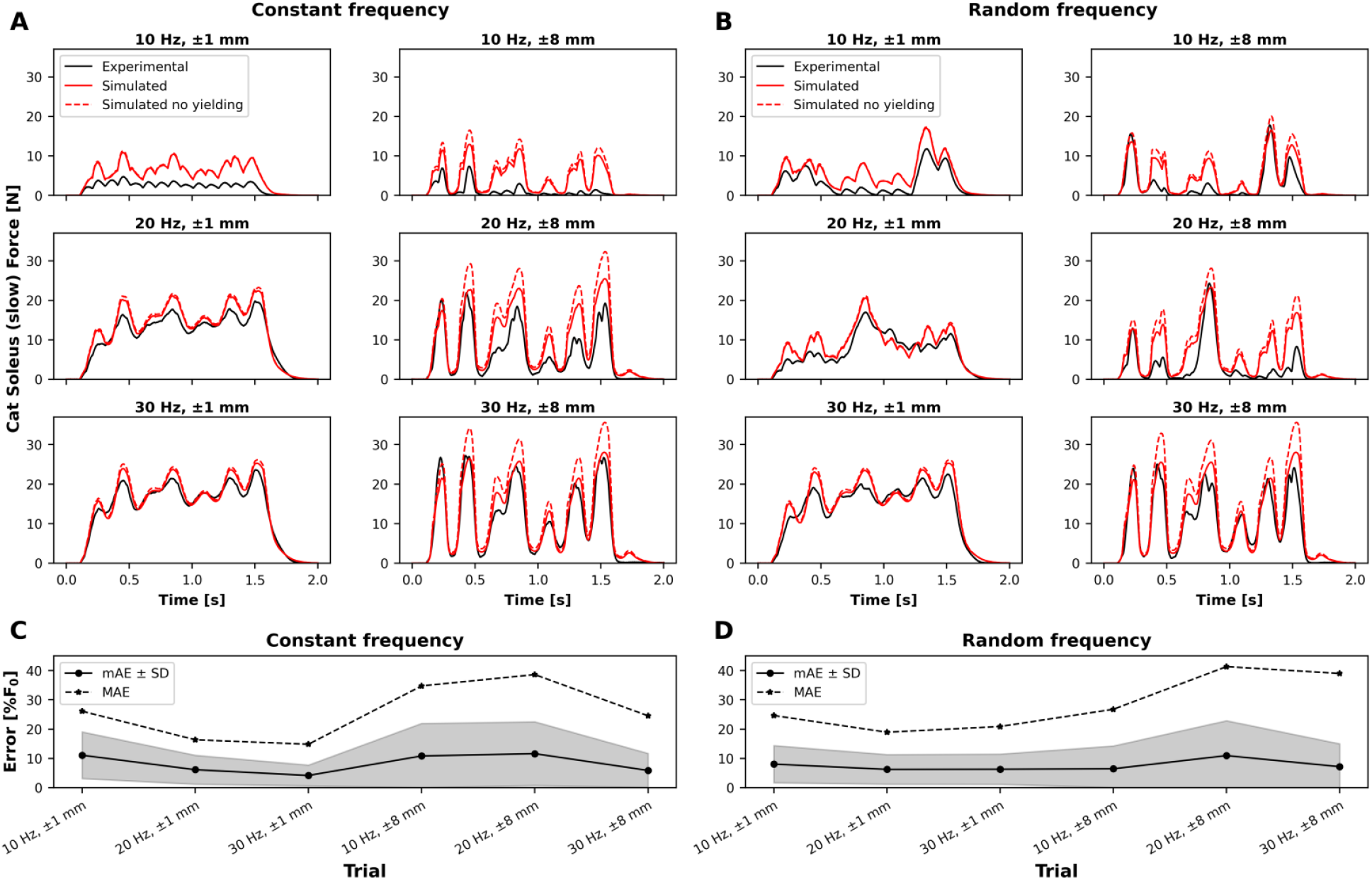
Experimental and simulated force responses of the cat soleus muscle-tendon unit (slow whole-muscle scale; 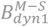 benchmarks) subjected to length changes with maximum amplitude of 1 mm and 8 mm. Simulations were performed under constant (A) and random (B) stimulation frequency (*f* ^*s*^) with mean values of 10 Hz (first row), 20 Hz (second row), and 30 Hz (third row) [14]. Mean absolute error (mAE, solid black line) ± standard deviation (SD, grey shaded area) and maximum absolute error (MAE, dashed black line) for the 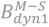 benchmarks are shown for constant-*f* ^*s*^ (C) and random-*f* ^*s*^ (D) trials, with yielding included in the simulations.

To further examine the FL and FV dependencies of the CE under maximal stimulation conditions, the 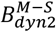 benchmarks were used (Fig 9A). Considering the 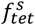 applied, the FL and FV relationships of the CE were tested when minimally affected by the NE active state, which was equal to 1 for the entire simulation time except the initial force ramp. The model reproduced the dynamic behaviour of the rat soleus muscle-tendon unit, achieving a mean mAE of 4.9 ± 2.5% and a MAE between 10.1% and 42.5% (Table 4). As shown in Fig 9B, the accuracy decreased progressively with increasing amplitude of length change, from mAE = 1.7 ± 1.5% (MAE = 10.1%) when the maximum imposed length variation was ± 0.05 mm, to mAE = 8.7 ± 7.3% (MAE = 42.5%) when the maximum amplitude reached ± 2.0 mm.

**Fig 9.**
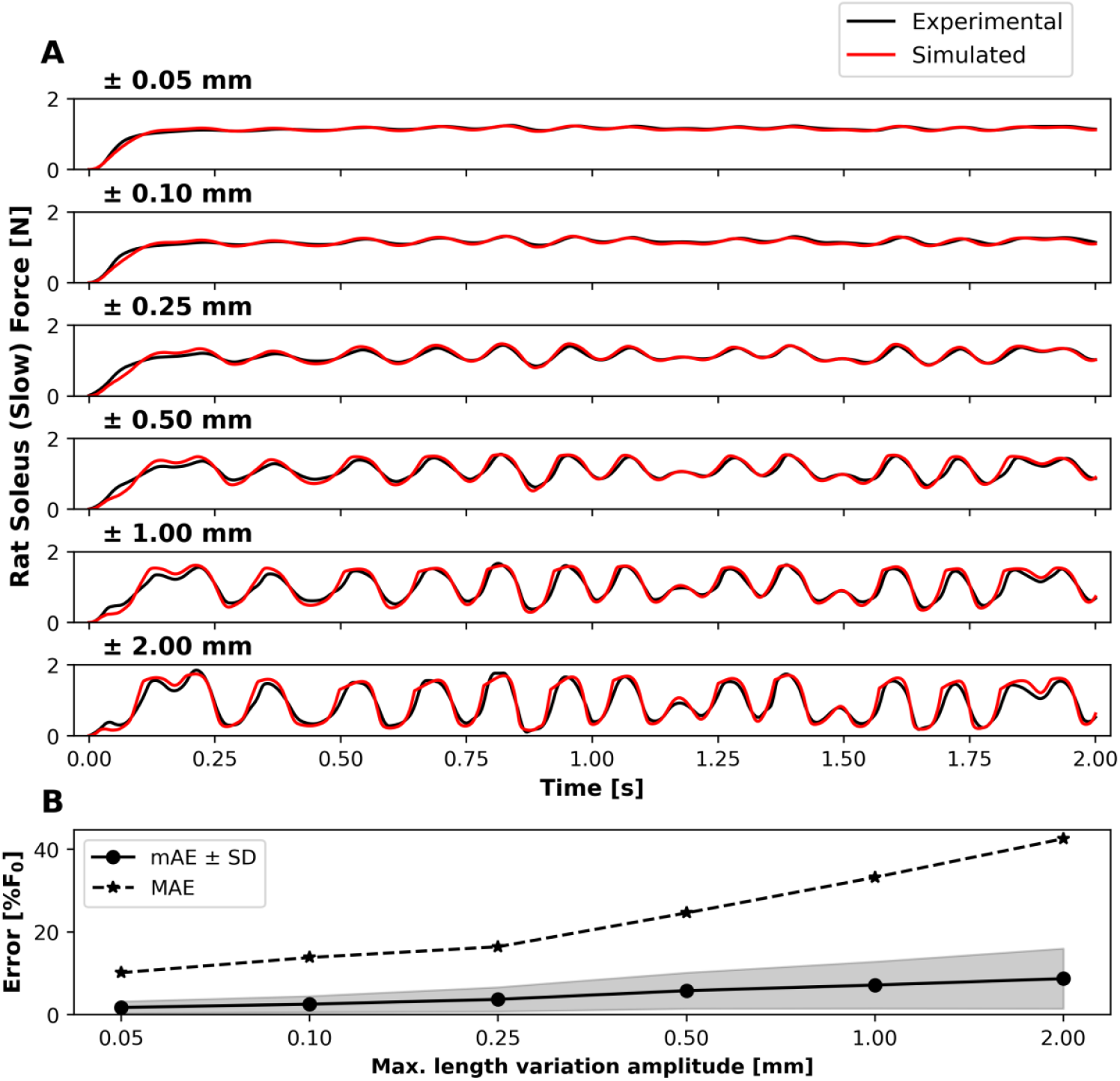
(A) Experimental and simulated force responses of the rat soleus muscle-tendon unit (slow whole-muscle scale; 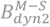 benchmarks) subjected to increasing length changes with maximum amplitudes of 0.05, 0.1, 0.25, 0.5, 1.0 and 2.0 mm (from the first to the sixth row) under maximal stimulation of 70 Hz [23]. (B) Mean absolute error (mAE, solid black line) ± standard deviation (SD, grey shaded area) and maximum absolute error (MAE, dashed black line) for the 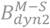 benchmarks.

### 3.3 Fast muscle benchmarks

#### 3.3.1 Isometric fast muscle benchmarks

The 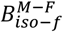 benchmarks (Fig 10A) were used to assess the model’s force-frequency relationship at the fast-type muscle scale. The model reproduced the experimental forces of the rat EDL muscle-tendon unit with a mean mAE = 6.9 ± 2.8% and 7.9% ≤ MAEs ≤ 28.4% (Fig 10C, Table S6 in Supplementary Materials S7). However, the experimental force-frequency relationship reached saturation at lower stimulation frequencies compared to the model. In the experimental data, at 50 Hz the muscle already reached at least 80% of its maximum force (≈ 1.3 N), whereas the model required at least 80 Hz to achieve comparable values (Fig 10A, 2^nd^ and 3^rd^ columns). Accordingly, the largest errors were observed at intermediate *f* ^*s*^ values, with a maximum mAE of 10.5 ± 8.8% at 50 Hz (Fig 10C), which progressively decreased with increasing *f* ^*s*^ to a minimum of 4.6 ± 4.8% at 100 Hz (excluding the 120 Hz trial used for calibration). The model accurately reproduced the experimental twitch response (mAE = 1.51%), although with a longer decay phase (*τ*_*decay*_ = 43 *ms*) (Table 5). Similarly, the fused tetanic response at 100 Hz exhibited a longer rising phase (*τ*_*rise*_ = 19 *ms*) (Table 5).

**Fig 10.**
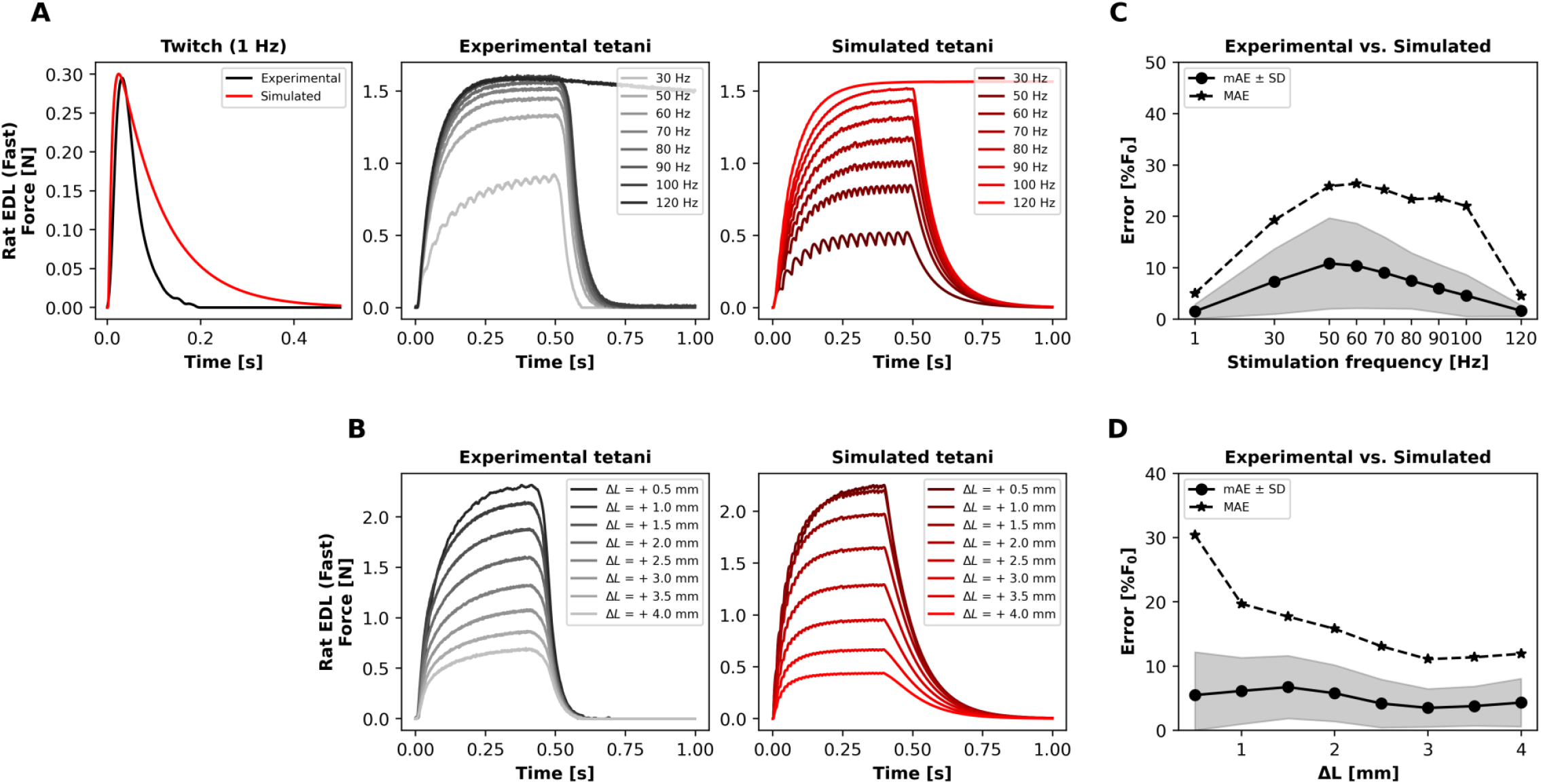
(A) Experimental and simulated isometric force responses of the rat extensor digitorum longus (EDL) muscle-tendon unit (fast whole-muscle scale; 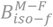 benchmarks). The first column shows the twitch response elicited at stimulation frequency *f* ^*s*^ = 1 Hz while the muscle–tendon unit was held at length 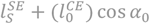. The second and third columns show experimental and simulated responses, respectively, for stimulation frequencies ranging from 30 to 120 Hz while the muscle–tendon unit was held at length 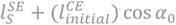. (B) Experimental (second column) and simulated (third column) isometric force responses (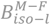 benchmarks). Simulations were performed at constant *f* ^*s*^ = 80 Hz while the muscle–tendon unit was held at a length of 30 *mm* + Δ*l*, with Δ*l* varying from 0.5 to 4.0 mm in 0.5 mm increments. (C-D) Mean absolute error (mAE, solid black line) ± standard deviation (SD, grey shaded area) and maximum absolute error (MAE, dashed black line) for the 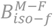 (C) and 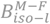 (D) benchmarks.

The 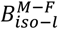 benchmarks (Fig 10B), performed on the same EDL muscle, were used to assess the FL relationship of the model. Across all length conditions, the model reproduced the experimental force responses with mAE ranging from 7.6% at Δ*l* = +1.5 *mm* to 3.6% at Δ*l* = +3.5 *mm* (Fig 10D). Similarly, the MAE decreased progressively from 30.4% at Δ*l* = +0.5 *mm* to 11.2% at Δ*l* = +4.0 *mm* (Table 4), with the highest errors observed at shorter length increments (Δ*l* ≤ 1.5 *mm*, see Fig 10D). Across both 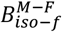 and 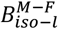 benchmarks, the simulated FI was slightly lower than the experimental FI, with a mean ΔFI of 0.2 ± 0.4 (Table 5). Notably, sag was not observable in either benchmark, as the time instant *t*_*p*_ could not be identified within the active force trace, and therefore was not included in these simulations.

#### 3.3.2 Dynamic fast muscle benchmarks

The 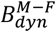 benchmarks (Fig 11) were used to validate the FV relationship of the model under sub-maximal and maximal stimulation conditions at the fast-type muscle scale. Under maximal stimulation (120 Hz), the model reproduced the experimental cat CF force satisfactorily during lengthening (mean mAE = 13.2 ± 4.7%) and accurately (mean mAE = 2.4 ± 2.0%) during shortening (Table S6, Supplementary Materials S7). In contrast, the model accuracy was reduced under sub-maximal stimulation. Across these trials (*f* ^*s*^ between 20 and 60 Hz), the simulations yielded 22.9 ≤ MAE ≤ 46.8%, with the largest MAE reached in the 40 Hz trial at 0.8*l*^*CE*^ for both shortening (46.8%) and lengthening (41.4%) (Table S6, Supplementary Materials S7).

**Fig 11.**
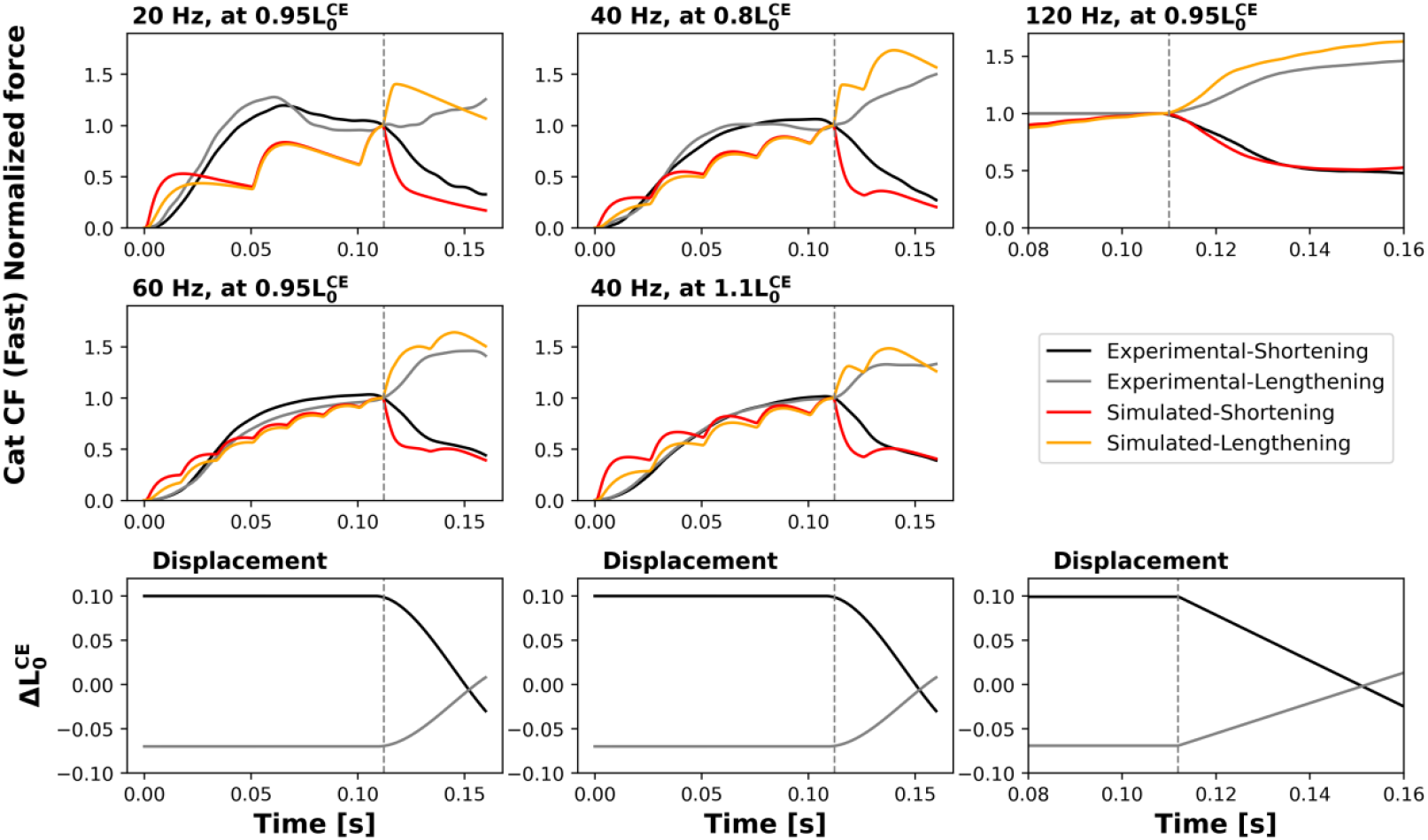
Experimental and simulated force responses of the cat caudofemoralis (CF) muscle (fast whole-muscle scale; 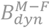 benchmarks). Unfused tetani (20, 40, and 60 Hz; first and second columns) and fused tetani (120 Hz; third column) are shown in response to prescribed shortening and lengthening displacements (third row) at different contractile element lengths expressed as fractions of the optimal fibre length 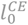 (0.95, 0.8 and 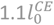). The experimental traces were manually digitised from [45].

## 4 Discussion

We aimed to determine whether a single Hill-type neuromuscular actuator, based on the MN-driven framework of [18] with addition of experimentally based excitation-activation dynamics, could accurately reproduce force traces recorded from experimental setups from the literature and collected *ad hoc* for this study. By validating the model against a comprehensive set of experimental animal benchmarks, including 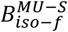 and 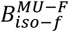 in isometric conditions for slow and fast MUs, respectively, and *B*^*M*−*S*^ for slow muscle and *B*^*M*−*F*^ for fast muscle in both isometric and dynamic conditions, we provide the first systematic assessment of a Hill-type neuromuscular actuator across scales, fibre types, stimulation regimes, and contraction modes. Overall, the results indicate that the proposed formulation could reproduce the main features of force production across these different conditions, while also highlighting specific limitations related to the modelling of sub-maximal activation, force-length interactions, and the representation of muscle activation with a single actuator.

The excitation and activation dynamics included in the presented Hill-type actuator differ from previous models in several ways. A first distinction pertains to how the active state is generated from the neural inputs. Most previous studies used Hill-type models based on either simplified first-order activation dynamics [34, 63], derived the active state *a posteriori* by inverting contraction dynamics from experimentally measured force [14, 15], or assumed it to be maximal throughout the entire simulation [23]. In contrast, our model predicts the muscle’s active state from the discharge instants via physiologically based calcium kinetics, inspired by the first conceptual formulation proposed by Hatze [20, 27]. Specifically, the excitation-activation dynamics is described by three ODEs (two 2nd-order and one 1st-order equation, see Supplementary Materials S1), which are based on experimental evidence and incorporate calcium transients derived from both slow and fast fibres. These features make the present formulation substantially less complex than alternative chemical approaches, such as [64], which used six 1st-order ODEs to describe the calcium-troponin binding process, or [65], which used 10 states to explicitly represent the excitation-contraction coupling. On the other hand, it remains considerably more flexible than simplified mathematical two-1st-order-ODEs approaches such as [66], which was designed for use in isometric conditions only. A second important distinction is that calcium dynamics depends non-linearly on muscle length at the level of the sarcoplasmic [Ca^2+^] transient, in accordance with experimental observations discussed in the present study (see Supplementary Materials S1 for more details). This contrasts with the only other MU-level-MN-experimentally-driven approach available in the literature [21], which assumes a non-physiological linear summation of MU twitches during muscle contraction [67, 68] and lacks any length-dependent modulation of the excitation-activation processes. To date, the only calcium-kinetics-based model that incorporated this length dependence is the AP-driven formulation developed by [22], which simulated force production using a single actuator representing the entire muscle, without distinction between slow and fast fibres. However, in that model, length dependence was introduced *a posteriori* based on observed force outputs rather than being based on direct experimental measurements of [Ca^2+^] dynamics. Moreover, this effect was implemented through modulation of the calcium-troponin binding rate constant, a downstream process relative to the calcium transient itself, where additional length-dependent mechanisms may superimpose and limit physiological interpretability [18]. A third relevant aspect concerns the calibration of the model’s parameters. The active state and the FV relationship represent the only stages of the proposed framework requiring parameter calibration. Notably, a total of only four parameters (*c*_*max*_, *k*_1_, *k*_2_ in Eq. 2 and a_*f*_ in Eq. 3 and Eq. 4) were calibrated (assuming that *F*_0_ is experimentally measured), compared to the twenty-three and twenty-eight parameters required by the models of [22] and [69], respectively, both including a mechanistic description of the excitation-activation dynamics. Most other models in the literature, in their excitation-activation dynamics did not explicitly represent the physiological processes linking neural input to muscle activation yet still required the calibration of a larger number of parameters. For example, neural impulse-driven approaches such as [66] relied on five calibrated parameters, while more popular EMG-driven models [39, 70] required eighteen.

The accuracy of the active state computed by the proposed excitation and activation dynamics can be evaluated indirectly through the force-frequency relationship under isometric conditions, as suggested by [8]. The mean mAEs across the 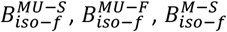 and 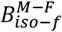 benchmarks (Table 4) were always below 11%, suggesting that the activation dynamics, based on mammalian data [28-32], can overall be extrapolated across different muscle scales, supporting the validity of the fibre-to-MU-to-muscle scaling [11] for this modelling stage. However, it is worth noting that both at the slow- and fast-muscle scales, the experimental force approached its fused-tetanic level at lower *f* ^*s*^ than in the simulations (Fig 6A and Fig 10A), particularly at intermediate *f* ^*s*^ (e.g. 10 Hz in the cat soleus or 50 Hz in the rat EDL). This suggests the presence of additional non-linearities in the excitation-activation cascade, or fibre-type-specific recruitment and saturation mechanisms, that may be missing in the current model. Nevertheless, even under identical experimental conditions, force responses may exhibit an inherent degree of variability. This is observed both at the muscle scale, as reflected by the discrepancies between the 10 - 20 Hz trials in Fig 6A and Fig 7A (obtained from the same cat soleus preparation in [14]), and at the MU level, as reported in [5], where force variations were a consequence of fatigue or potentiation phenomena.

The coupling between activation and contraction dynamics can be observed in isometric and dynamic trials. Looking at the force-length relationship under isometric conditions can help clarify. The error trend of the 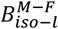 benchmarks at constant *f* ^*s*^, with mAE always below 8% (Fig 10D) indicate that the model adequately reproduced the effect of length variation on force output at the fast muscle scale, with improving accuracy as the muscle is stretched further. In contrast, in the 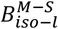 benchmarks the model accuracy progressively decreased at higher frequencies, reaching mAE = 24.7% for Δ*l* = 0 *mm* at *f* ^*s*^ = 40 Hz (Fig 7B). These results were due to the model being unable to capture how, in the experimental trials, the increasing stimulation compensated for the change in force-generating capacity due to the shift along the FL curve, resulting in very similar force responses across different lengths, especially for *f* ^*s*^ > 20 Hz (Fig 7A). It is plausible that this behaviour could be reproduced by a more complex interaction between the FL relationship and the non-linear dependency of calcium kinetics on muscle length. Nevertheless, the model demonstrated robust performance in the dynamic 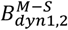 benchmarks, where the interaction between the activation and contraction dynamics is maximum, achieving an accuracy comparable to that reported by previously employed Hill-type formulations [14, 15, 22, 23]. Specifically, under maximal activation conditions in 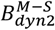, the model achieved mAE values ranging from 1.7% to 8.7% (Table 4), demonstrating accuracy comparable to that obtained with the damped equilibrium model, and consistently outperforming the rigid-tendon model, both reported by [15]. In the sub-maximal stimulation dynamic benchmarks 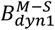, characterised by substantial modulation of the active state and known to be particularly challenging for standard Hill-type models [9, 13, 38], the present formulation maintained good predictive accuracy across all tested conditions, with 4.1 ≤ mAE ≤ 11.6 % when yielding was considered (Table 4). Notably, for the 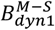 trial with the largest amplitude length excursions (± 8 mm), where traditional Hill-based approaches frequently exhibit considerable degradation in performance, the model preserved mAE values between 7.1% and 11.6% (Table S4 in Supplementary Materials S6), comparable to [15], where the active state was computed from the experimental force, rather than assigned as input. As previously discussed, this improvement was primarily achieved through the incorporation of yielding into the contraction dynamics of the model (Fig 8A, Table 4). A similar accuracy in the same 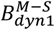 trials was previously obtained by the AP-driven model developed by [22], but only after introducing an *a posteriori* compensation that altered the activation dynamics to minimise the prediction error. Here, on the other hand, the yielding compensation was modelled phenomenologically based on experimental observations [45].

In this study, differently from classical Hill-type muscle models, the activation and contraction dynamics coupling extends also to the force-velocity formulation. Standard approaches [8, 63] typically assumed a decoupled activation and contraction dynamics, whereby force was computed as the product of the active state and independent *f*_*FL*_ and *f*_*FV*_. In contrast, the FV relationship in Eq. 3 and Eq. 4 explicitly incorporates non-linear dependencies on muscle length and activation through the modulation of the maximum contraction velocity *K*, including both force-velocity-length (FVL) and force-velocity-activation (FVA) interactions. Such properties remain uncommon in the Hill-model literature, despite their physiological evidence [9]. Indeed, a recent systematic review on Hill-type models [9] reported that only approximately one third of the analysed studies included length or activation dependencies in their FV formulation. However, the adoption of this FV relationship resulted into a worse predictive accuracy of the model in the 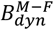 compared to the 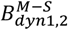 benchmarks (Table 4). This likely reflected a mismatch in the temporal structure of the activation input. The experimental cat CF data were generated through asynchronous stimulation of multiple ventral root bundles, that produced smooth force traces even at low *f* ^*s*^, and which *f* ^*s*^ patterns were unknown and therefore not reproducible in a simulation. On the other hand, our model relies on synchronous stimulation trains inherent to its single-actuator architecture. This clearly led to amplified force oscillations and accentuated deviations (Fig 11), particularly for sub-maximal activation. Consequently, the limited predictive accuracy observed in these dynamic conditions can reasonably be interpreted as the result of a scale mismatch between the model’s input, trying to closely represent the experimental conditions, and the simplified representation of the muscle as a single actuator, incapable of replicating the desynchronised recruitment and firing patterns characteristic of MU populations stimulated via ventral root bundles.

Notably, this is the first model since the Virtual Muscle formulation of [71] that incorporated both the time-dependent mechanisms of yielding (in slow fibres) and sag (in fast fibres). Yielding was modelled as being solely dependent on contraction velocity at sub-maximal *f* ^*s*^, which, consistent with experimental evidence [46], ensured that it does not substantially affect simulations under isometric or quasi-isometric conditions and, as discussed above, consistently improved the model predictions. The sag implementation, on the other hand, was modified from [47] by restricting its action to sub-maximal stimulation conditions and by introducing as an input the time instant *t*_*p*_. This makes the formulation more robust, as *t*_*p*_ is a measurable quantity rather than an arbitrary threshold, such as the *f*_0.5_ criterion originally adopted by [47], but also limits the applicability of the model for standard musculoskeletal simulations, in which *t*_*p*_ is not available. Unlike yielding, the inclusion of sag did not result in a significant improvement in predictive accuracy across the 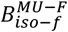 benchmarks. This likely reflects a strong sensitivity of the sag response to the calibration of the phenomenological parameters *T*_*S*_, a_*S*1_, and a_*S*2_ (Eq. 6 and Eq. 7), which determine both the temporal evolution and amplitude of the force decay. Moreover, as observed in the experimental unfused tetanic contractions of the cat and rat MG MUs (Fig 5), the magnitude and shape of sag differed substantially between the two animal species. This suggests that sag-related parameters may require dedicated calibration for each experimental preparation. While the effect of sag on overall predictive accuracy was modest, reducing the mean MAE from 25.6% to 24.5%, yielding consistently improved model predictions, reducing the mean MAE from 37.1% to 27.2% (Table 4), These results support the inclusion of yielding when modelling slow fibres under dynamic conditions, whereas the role of sag may require further investigation using larger datasets including different muscles and species. This study is not without limitations that warrant consideration. A first limitation relates to the scarcity of data reporting experimental [Ca^2+^] transients across fibre types, sarcomere lengths, and temperatures, which prevents a direct assessment of whether the highest errors obtained across the *B*_*iso*−*f*_ benchmarks originate primarily from the calcium dynamics or from the link between calcium and active state. In particular, the lack of [Ca^2+^] data for slow-fibres at a physiological temperature [30, 32] limits the ability to assess the validity of the assumption made regarding the [Ca^2+^] temperature scaling for slow-type kinetics (Supplementary Materials S1). The lack of extensive data in this domain also prevents a deeper investigation into the appropriate modelling stage at which time-dependent phenomena such as yielding and sag should be introduced, which is currently unclear. In the present study, they were applied as generic scaling factors acting directly on the active state. However, it is plausible that they originate from more distinct intra-sarcomere mechanisms. Specifically, yielding is likely driven by an increase in the cross-bridge detachment rate during active lengthening [45], while sag has been hypothesised to result from an increased rate of sarcoplasmic calcium removal [47]. Incorporating these phenomena directly into the [Ca^2+^] level (Eq. 1) and active state (Eq. 2) ODEs would allow a more mechanistically accurate representation. Secondly, the present Hill-type formulation overlooks the active viscoelasticity of the CE. Consequently, the model cannot capture the short-range stiffness and rapid force transients that occur during small length perturbations [72]. However, this simplification is consistent with the primary aim of this work, which was the development and validation of the excitation-activation dynamics and its interactions with the contraction dynamics, rather than the detailed representation of muscle impedance. Moreover, under the physiological conditions included in our benchmarks, advanced formulations that include viscoelastic cross-bridges have been shown to rapidly converge to the force outputs predicted by traditional Hill-type dynamics over time [73], further justifying our decision to omit this feature. A third limitation concerns the limited benchmark data currently available at the MU scale. To the best of the authors’ knowledge, no dynamic MU force data are available in the literature, restricting the validation of the model at the MU level to isometric conditions only. Moreover, the available datasets cover only a narrow range of fibre-type properties extrapolated from FR units and, in the case of the cat MG, include the sag effect in only a single sub-maximal stimulation trial. Similarly, the muscle benchmarks are based on specific animal preparations (cat soleus, rat soleus, rat EDL, cat CF) and experimental protocols that generally do not cover the full spectrum of physiological loading and activation patterns encountered *in vivo*. These constraints, together with differences in temperature, fibre-type composition and tendon architecture between species, introduce multiple sources of uncertainty and prevent a confident extrapolation of this study findings to other muscles or to human conditions. On the other hand, this limitation also clarifies the value of the presented comprehensive benchmarks, which are available online at https://github.com/andreasgarzi/MuscleModelBenchmarks together with an implementation of the presented model, for the computational biomechanics field and for the assessment of other neuromusculoskeletal Hill-type actuators used at multiple scales (e.g. [18, 21, 27]).

The findings and limitations of this study suggest several directions for future work. First, they highlight the need for systematic experimental datasets of dynamic MU contractions and [Ca^2+^] transients across multiple fibre types, temperatures, and sarcomere lengths within the field of computational muscle modelling. Such data would allow further refinement of the excitation-activation formulation and a more comprehensive assessment of Hill-type actuators operating at the MU scale. For the Hill-type neuromuscular actuator presented in this study, future developments will consist of combining single-actuators into a population of MUs at the muscle scale, incorporating realistic MU recruitment and firing strategies, as previously implemented in [18]. This, coupled with inputs from experimentally derived MN discharge patterns [74-76], will enable the study of how MU pool organisation and asynchronous activation shape the macroscopic force response under different modes of human muscle contractions, extending previous work from [18, 77]. Such developments would further consolidate the model validated in this work as a versatile tool for neuromuscular research and for the interpretation of neural control strategies in both healthy and pathological conditions.

## 5 Conclusion

This work provides the first systematic validation of a multiscale Hill-type neuromuscular actuator across physiological scales, fibre types, and contraction conditions using a structured set of experimental benchmarks. The proposed ODE-based excitation-activation framework, based on calcium kinetics and coupled to a three-element Hill-type structure, constitutes a physiologically interpretable model to reproduce the main features of muscle force production under the simulated experimental conditions. Importantly, the model generates force directly from discharge times as the sole control input, which cannot be achieved with classical Hill-type formulations. Across benchmarks, mean errors were typically below 15% *F*_0_, although larger errors were observed in specific trials, particularly under submaximal dynamic conditions. This performance was achieved with only four calibrated parameters of the active-state and force-velocity relationships. Moreover, the inclusion of physiologically based excitation-activation dynamics, together with yielding and sag as fibre-type-specific effects, improved the ability of the model to reproduce force responses under submaximal activation conditions. Therefore, the present study represents an important step toward the systematic validation and development of Hill-type neuromuscular actuators, including formulations with MU-level resolution.

## Credit authorship contribution statement

**Andrea Sgarzi:** Conceptualisation, Methodology, Software, Data Curation, Validation, Formal analysis, Investigation, Visualisation, Writing - original draft.

**Arnault H. Caillet**: Conceptualisation, Methodology, Writing - Review & Editing.

**Matthew Millard**: Methodology, Investigation, Writing - Review & Editing.

**Sven Weidner**: Methodology, Investigation, Writing - Review & Editing.

**Nicos Haralabidis**: Methodology, Writing - Review & Editing

**Théo Meranger**: Conceptualisation, Methodology, Writing - Review & Editing.

**Bart Bolsterlee**: Supervision, Writing - Review & Editing.

**Dario Farina**: Conceptualisation, Supervision, Writing - Review & Editing.

**Nigel H. Lovell**: Supervision, Writing - Review & Editing.

**Luca Modenese**: Conceptualisation, Methodology, Resources, Visualisation, Supervision, Writing - original draft, Writing - Review & Editing, Funding acquisition.

## Acknowledgements

This research was supported by a Scientia Fellowship granted by the University of New South Wales to Luca Modenese. Matthew Millard and Sven Weidner gratefully acknowledge financial support from the Deutsche Forschungsgemeinschaft through project 540349998.

